# Fishing for survival: importance of shark fisheries for the livelihoods of coastal communities in Western Ghana

**DOI:** 10.1101/2021.01.18.427106

**Authors:** Issah Seidu, Lawrence K. Brobbey, Emmanuel Danquah, Samuel K. Oppong, David van Beuningen, Moro Seidu, Nicholas K. Dulvy

## Abstract

Small-scale shark fisheries support the livelihoods of a large number of coastal communities in developing countries. Shark meat comprises a cheap source of protein and is traded locally in many parts in developing countries, while the skins, oil, and fins are exported to the international market. This study addresses a gap in literature regarding the importance of elasmobranchs to key shark-fishing communities and the degree to which trade in shark products (meat and fins) vary in time and among fishing communities in Ghana. We interviewed 85 fishers and traders involved in shark fisheries in Axim, Dixcove, and Shama communities using semi-structured questionnaires. Fishing was the primary source of income and accounted for 58.5% of the total household income of respondents. Other important economic activities were fish processing (16.0%), fish retailing (13.3%), and small businesses (2.5%). One-third and often two-thirds of respondents generated between 80-100% of their income from shark fisheries: Axim (65%), Dixcove (68%), and Shama (35%). Shark meat consumption was common among fishers and traders and represents a substantial source of protein in the diet of the study communities. Hammerhead sharks (*Sphyrna* spp) and Bull Shark (*Carcharhinus leucas*) have the most valuable fins and meat. Further, 75% and 95% of fishers and traders, respectively, see fishing and trading of shark meat as their last safety-net and, therefore, tend to be satisfied with their jobs. Non-fishing related livelihood streams including small businesses and transportation were the major fallback activities both fishers and traders preferred to rely on if there is a ban on the exploitation of sharks in Ghana. Overexploitation of these species will compromise food ecosystem functionality and security. Thus, any shark management strategy needs to urgently restraint mortality to sustainable levels, which, in the short-term, must take into consideration the preferred livelihood fallback options outlined by fishers and traders, and implement them to ensure the long-term benefits of the intervention.

## 1. Introduction

The economies of developing countries heavily depend on natural resources-based livelihood strategies, with fisheries contributing significantly to the economy (BNP, 2008). For instance, the United Nations Food and Agriculture Organization (FAO) and the World Fish Centre-Big Numbers Project (BNP) (2008) estimated that between 93 and 97 million rural households in developing countries worldwide are involved in fishing and its related activities. An estimated 12.3 million people in Africa are actively involved in fishing as either fishers, processors or traders (FAO, 2014). The fisheries sector accounts for more than US$ 24 billion or 1.3% of the joined GDP of all African nations and also supports 30% of Africa‟s nutrition and food security (FAO, 2014). In Ghana, the fishery sector directly or indirectly employs 2.6 million people, comprising 10% of the total population (Nunoo et al., 2015). The fishery sector is estimated to support about 3% of Ghana‟s GDP and 5% of Agricultural GDP (Nyemah et al., 2017). The sector also contributes 60% of animal protein consumed in Ghana with average per-capita consumption between 20 and 25 kg per annum (Nunoo et al., 2015).

The inshore artisanal fishery is a major source of Ghanaian fish production, accounting for 80% of the total annual fish supply (MoFAD, 2017). The main aim of artisanal fishers is to continue to supply fish to meet their household needs and for domestic markets. The artisanal fishery operates actively in over 300 landing sites in several coastal communities along the 550 km coastline of Ghana and supports approximately 124,000 fishers (Amador et al., 2006; CRC, 2013). A number of coastal communities in Ghana engage in subsistence farming, small trade, artisan works, factory work, mining, and sand mining as their livelihood strategies and these are mostly undertaken concurrently with fishing-related activities (Mensah & Antwi, 2002). However, recent studies have found that artisanal fisheries are the most important livelihood strategies for coastal communities in Ghana. For example, Asiedu and Nunoo (2013) report that between 80% and 97.7% of fishers from the studied population depend on fishing as their sole occupation in the Ghanaian coastal communities of Small London, Kpong, Ahwiam, and Elmina. Similarly, Asiedu et al. (2013) report that artisanal fishing contributes 80% and 85% to the total household income of fishers in the Ahwiam and Elmina communities, respectively.

Livelihoods of many artisanal fishing communities in sub-Saharan Africa are under threat, as the region continues to put increasing pressure on fish resources for sustenance and income, which poses many challenges for long term sustainability of the resources for food security (Andrew et al., 2007; Béné et al., 2005). This has caused many fishers to resort to diverse strategies to maintain or improve their livelihoods. The adopted strategies include geographical mobility, utilizing different methods of fishing, fishing in different locations, and adjustment from specialist to generalist fishing operations (Allison & Ellis, 2001; Smith & Mckelvey, 1986). The latter is the most common strategy among Ghanaian artisanal fishers in an effort to maintain high levels of catch in the wake of continuous declines of their formerly preferred teleost stocks (MoFAD, 2015). These generalist fishers invariably use diverse fishing gears or modify their fishing operations to target bony fishes, marine invertebrates, and other vulnerable marine megafauna (I. Seidu, pers. obs.). Elasmobranchs (sharks, rays, and skates) are among the marine megafauna that are particularly susceptible to capture in diverse fishing gears and across a magnitude of fishing operations, with fisheries posing the greatest source globally to non-natural mortalities within this group (Bonfil, 2000; Dulvy et al., 2000; Dulvy et al., 2014).

Elasmobranchs typically have a relatively slow life history due to their large body size, late maturity, slow growth, and low fecundity, which results in low population growth rates (Pardo et al., 2016). These traits make them exceptionally vulnerable to overfishing and typically result in decreased chances of recovery from population decline (Barrowclift et al., 2017; Dulvy et al., 2000). Traditionally, elasmobranchs primarily constituted bycatch until the rise in international demand and prices for their products, particularly for fins in the mid-1980s, which incentivized many coastal communities to target sharks and rays (Clarke, 2004). Shark fins are now rated as one of the most expensive fish products worldwide resulting in some sharks and rays being the most valuable traded wildlife (McClenachan et al., 2016). Although there has not been a robust estimate of the number of people involved in small-scale elasmobranch fisheries worldwide, this activity has been recognized to support a large number of rural coastal community livelihoods in developing countries (Bonfil, 2000). Elasmobranch meat is traded locally in many parts in developing countries and can form a cheap source of protein for the people, for example, in Southern Brazil (Bornatowski et al., 2013).

Recent studies in the sub-Saharan African region have demonstrated shark fisheries as an important livelihood strategy for coastal communities (Barrowclift et al., 2017; Diop & Dossa, 2011; Gelber, 2018). For example, in the Sub-Regional Plan of Action for Sharks (SRPOA-Sharks) project, Diop & Dossa (2011) indicate that shark fishing provides an estimated 13,000 direct jobs to fishers, processors, and fish smokers in 2008 of, which 7% was generated by artisanal fishing in the Sub-Regional Fisheries Commission Zone. In Zanzibar, East Africa, Barrowclift et al. (2017) report that elasmobranchs contributed 41-60% of the total income of fishers who caught and sold sharks. They also report that 31% of merchants obtained between 61-80% of their income from selling elasmobranchs. Additionally, in Ghana, trade in shark fins is the main source of income for 80% of middlemen and 38% of canoe owners of the study population (Gelber, 2018). Although these studies provide initial data on trade in shark fins (Gelber, 2018) and meat as well as livelihood strategies of fishers (Barrowclift et al., 2017), the historical trade dynamics of fins and the local consumption pattern of shark meat have not been documented. Since most rural coastal communities depend on shark meat for their protein requirements, enquiries about consumption of shark meat is relevant for designing management strategies for the sustainable benefit of the rural communities. Further, comparatively, few studies have investigated the wellbeing and income satisfaction in artisanal fisheries, which employs over 90% of fishers globally (FAO, 2014; Purcell et al., 2016). Fishers‟ satisfaction may be influenced mainly by income and happiness, and none-monetary factors such as adventure and self-actualization (Pollnac & Poggie, 2006; Coulthard, 2011). Satisfaction impacts the health of fishers and the relationship between fishers and management institutions, and offers opportunity to target training and development programs for fisheries (Ruiz, 2012; Trimble & Johnson, 2013).

Additionally, many species of elasmobranch are considered threatened as a result of direct impact from target and bycatch fisheries worldwide (Dulvy et al., 2014; 2017) and such is the case of the shark fauna in Ghana. Most fishers in a previous study reported that shark species such as Hammerhead sharks (*Sphyrna* spp), Thresher sharks (*Alopias* spp), Lemon Shark (*Negaprion brevirostris*), and Mako sharks (*Isurus* spp) among others, have declined in recent years (Seidu et al., Submitted). With the current fishing pressure, it is likely most shark populations will continue to decline and some may even go extinct if effective management measures are not urgently instigated, as has occurred for sawfishes in the region (Dulvy et al., 2016; Fernandez-Carvalho et al., 2014). Following international concern over rapidly dwindling shark populations, several mitigation measures are advocated by biologists and conservationists. Spatial closures, such as marine protected areas (MPAs) and no take marine reserves as well as fishing bans are some of the fisheries management strategies that have been implemented to slow and reverse the effect of large-scale overfishing on shark populations (Ward-Paige et al., 2012). However, implementation of these measures has the potential to adversely impact regions with significant shark fisheries and stakeholders who directly depend on shark and other marine resources for their livelihoods. Thus, exploring potential effects of fishing bans or closures on fishers‟ behavior, especially fallback livelihood options they prefer to rely on, is not only important to ensure the welfare of fishing communities, but also to increase the chance of success of shark protection if management authorities are to implement such measures to mitigate shark decline in Ghana.

Ghana is among the major artisanal fishing nations, with long history of catching sharks since 1700s (Jorian, 1988). Since the late 1950s, shark landings have been increasingly erratic in Ghana, peaking in 1975 with 11,478 tons (FAO, 2017). In the last decade, the total reported shark catches fluctuated considerably. The catch peaked up to 10,000 tons in 2013 and dropped to 8,152 tons in 2015 (FAO, 2020). Since 2015, however, the catch estimate trends indicate a sharp decline in shark landings (FAO, 2020). The decline in shark catch corroborates a recent study on Local Ecological Knowledge of fishers that indicates a remarkable decline in shark catches since the 2010s, suggesting that sharks are overexploited in Ghanaian waters (Seidu et al., in review).

Given these considerations, this study aims to address a significant data gap on the economic impact of shark fisheries to fishers‟ and traders‟ livelihoods, and assesses the trade dynamics of shark products in major shark-fishing communities in Ghana. We specifically tackle the following questions: (i) what are the existing livelihood strategies and how is shark fishery contributing to artisanal fishers and traders‟ household income and sustenance? (ii) What are the fallback livelihood options available to fishers and traders in the advent of a ban on shark exploitation in Ghana? (iii) How have the sales prices of commercially important shark products (fins and meat) changed over time? and (iv) Are fishers and traders satisfied with their work and the income they obtained from shark products? The findings are essential in targeting policy interventions in livelihood enhancement, food security, and poverty reduction. In addition, understanding the livelihood strategies, fallback options, and wellbeing of primary actors in shark fisheries is necessary for planning management interventions for the sustainable utilization of sharks.

## 2. Methodology

We first described the analytical framework for the study. Second, we described the socio- economics, geography and historical catch trends of sharks in Ghana as well as the study areas for the study. Third, we described the data collection methods and finally, we detailed our data analysis methods.

### 2.1 Analytical framework

The analytical framework of the study was based on the Sustainable Livelihood Approach (SLA) (Figure 1). The SLA is founded on the premise that people require a range of strengths (here called “capitals”) to achieve positive livelihood outcomes (DFID, 2000; Ellis, 2000; Scoones, 1998). At the heart of the framework is the livelihood strategy, made up of natural resource-based (e.g., fishing, collection of aquatic resources, livestock farming) and non-natural resource-based (e.g., trading, artisanship, services) livelihoods in rural settings. The SLA identifies five types of capitals upon which the choice of a household to pursue a particular livelihood strategy is built, namely, natural, human, social, financial, and physical capital. Mediating institutions (e.g., policies, customs, taboos, and rules) determine access to various capitals and the choice of households to build a particular livelihood strategy. Mediating institutions further have direct influence on livelihood outcomes (e.g., whether fishers are able to achieve a feeling of inclusion and well-being) (DFID, 2000). The livelihood outcomes in turn have influence on these mediating institutions. External factors such as seasonality, critical trends, and shocks over which people have limited or no control, have influence on the wider availability of capitals and livelihoods of households (both livelihood strategy and outcome). Livelihood strategies undertaken by fishers may result in improved income levels, increased well-being, satisfaction, or sustainable use of fishery resources (livelihood outcomes). For instance, fishers may improve their income through diversifying their livelihood strategies, which will invariably reduce pressure on fishery resources resulting in sustainable use of the resources. Finally, the resulting livelihood outcome of fishers‟ households invariably influences their capital through investment in education of household members or financial savings.

**Figure 1.**
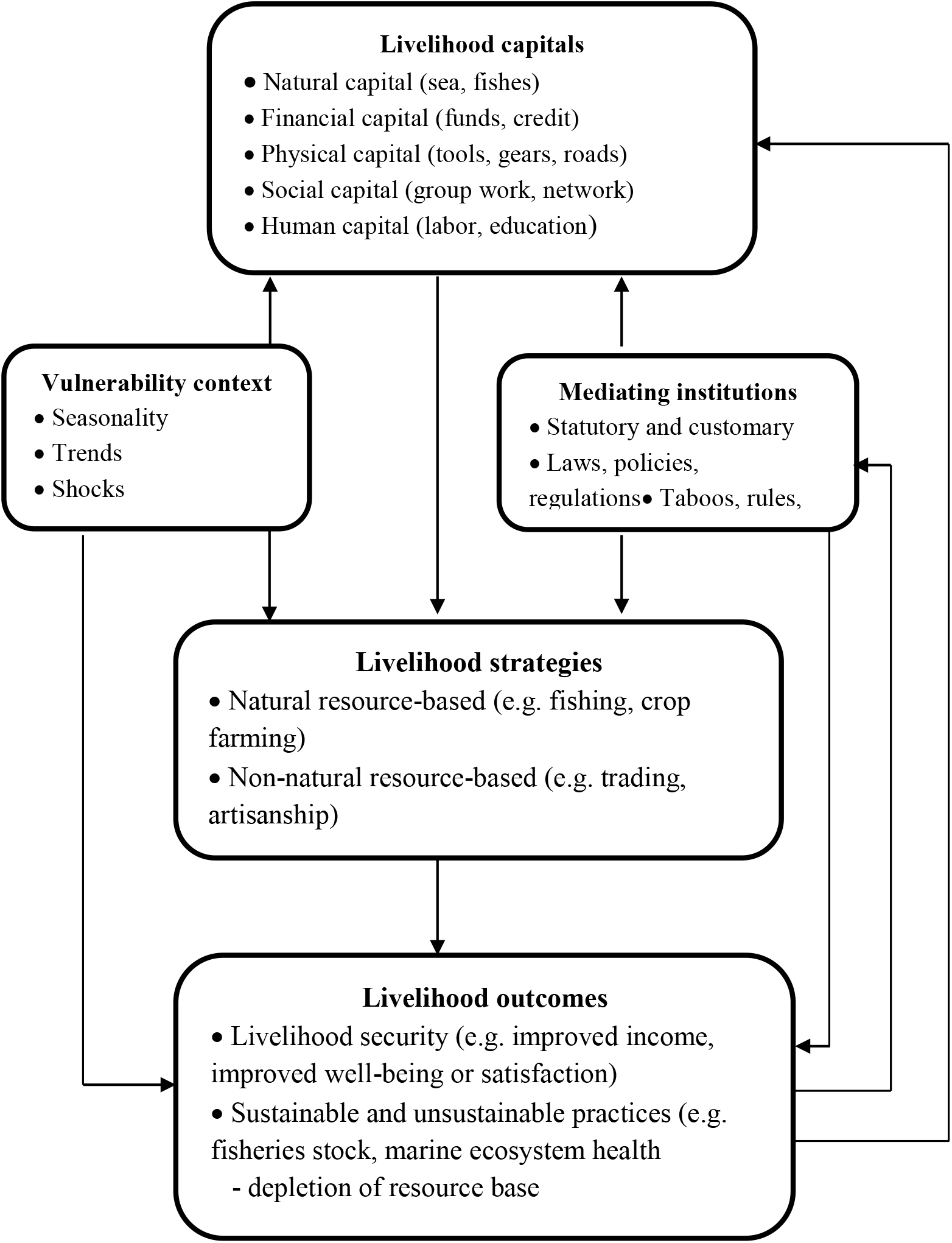
Analytical framework of sustainable livelihoods. Adapted from DFID, (2000), Ellis (2000) and Scoones (1998, 2015)

Information presented in this study mostly emphasizes the livelihood strategies and livelihood outcomes (income level and wellbeing) of the SLA analytical framework as highlighted in Figure 1. Other components of the framework including livelihood capitals, vulnerability, and mediating institutions have been addressed in our previous papers (Seidu et al., submitted).

### 2.2 Study Area

Ghana is a Western African nation bordered by the Burkina Faso to the north, Republic of Côte d’Ivoire (Ivory Coast) to the west, the Togolese Republic (Togo) to the east, and the Gulf of Guinea to the south. Ghana lies along the Gulf of Guinea (30 5’ W and 1010’ E and 40 35’N and 110 N) and has an area of about 239,000 km^2^. Ghana‟s coastline is approximately 550 km long with about 90 lagoons and associated wetlands. The coastal zone covers 6.5% of land area but is inhabited by a quarter of the population (deGraft-Johnson et al., 2010) and is split into three geomorphic units. The West Coast extends from the Ghana-Côte d‟Ivoire border to the Ankobra River estuary. The Central Coast from the Ankobra estuary to Tema has rocky headlands and sandbars enclosing coastal lagoons. The East Coast stretches from Tema to the Ghana-Togo border where the shoreline is sandy; this area is characterized by considerable erosion. Generally, the marine resources of Ghana encompass over 347 fish species, belonging to 82 taxonomic families (deGraft-Johnson et al., 2010).

The study was conducted in three coastal communities in the Western Region, along the West Coast, namely, Axim, Dixcove, and Shama (Figure 2, Table 1), which are the hotspots of shark fisheries in Ghana. The communities were chosen based on three major reasons – that is, fishing is exclusive to artisanal fishers; sharks form a significant catch and characterized with local shark fin trade; and fishers were willing to cooperate with the researchers for both landing and interview data. The Axim, Dixcove, and Shama communities fall within the Nzema East Municipality, Ahanta West, and Shama Districts respectively. Axim community, with a mean annual precipitation of 1,979 mm, exhibits the highest average rainfall pattern in Ghana (GSS, 2014), which favors crop farming activities. Dixcove and Shama have a mean annual rainfall of 1,700 mm and 1,820 mm, respectively.

**Figure 2.**
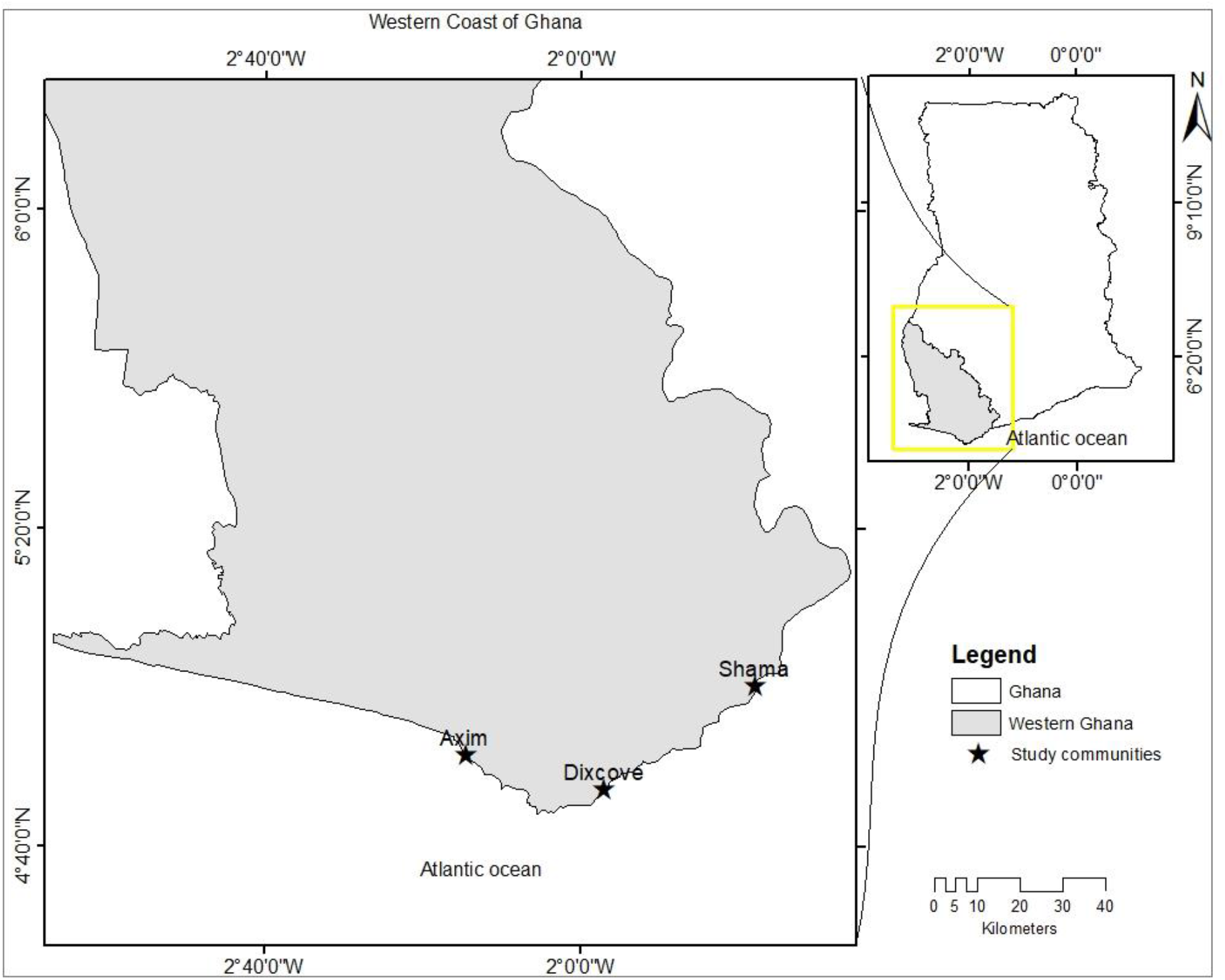
Map of Western Ghana showing the three study communities

**Table 1.**
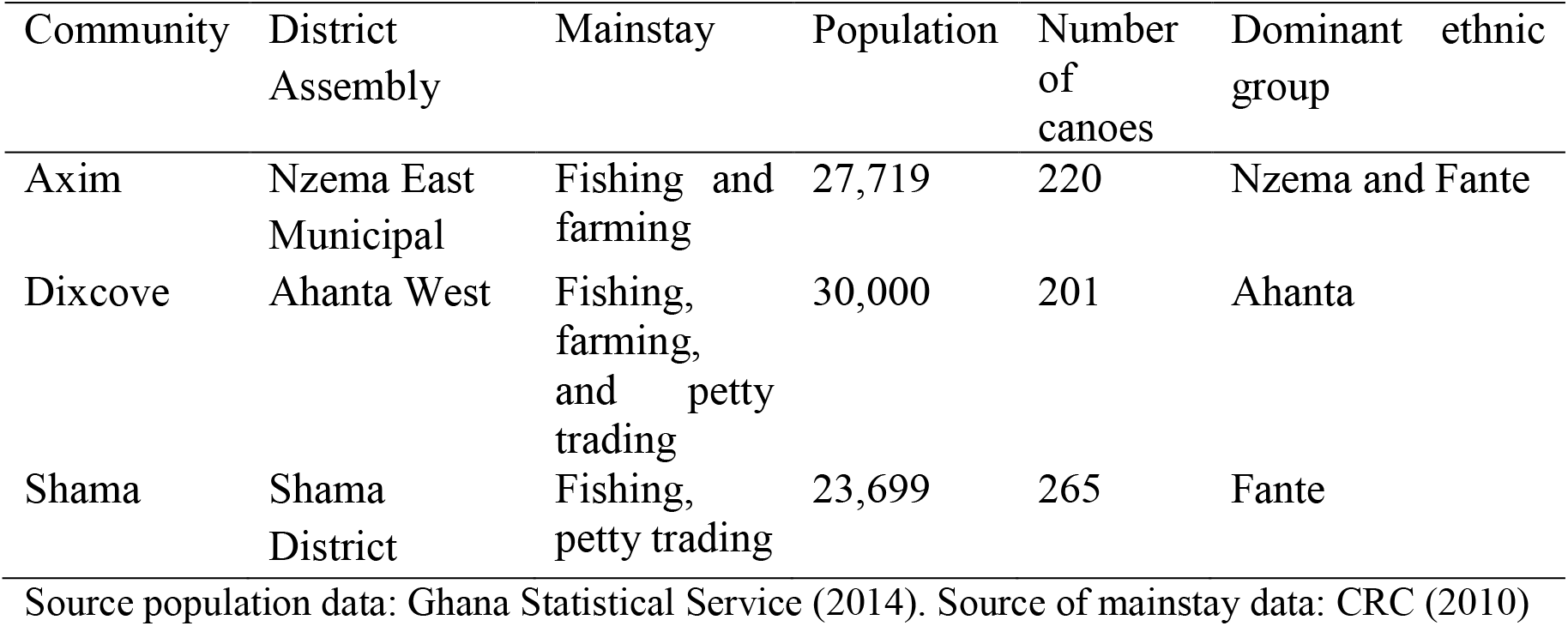
Characteristics of the study communities

### 2.3 Data collection

Data collection started in February 2020 and ended in August 2020. Data were collected using a semi-structured questionnaire that was designed to gather both qualitative and quantitative information. The questions for the interview were pre-tested for clarity in Shama and Axim with ten fishers (five from each community) in February 2020, as shark meat and fin trade are prevalent in these two communities. This gave an opportunity for us to make the necessary changes to reflect the local context of the study communities. Aside from administering all interview questions to respondents, we also specifically ask them to comment on how shark fishery affects their livelihood strategies and trade dynamics, and the influence of sharks on their income.

A non-probability convenience sampling approach (Alexander et al., 2017) was employed to select fishers and traders for the interview. The convenience sampling approach also referred to as availability sampling is based on the availability and willingness of respondents to participate in the interview (Naderifar et al., 2017; Newing, 2010). Thus, the number of respondents interviewed in each community depended on the availability and willingness of fishers and traders to participate in the interview. This sampling scheme was chosen because most fishers and traders were aware of the global controversies surrounding sharks and the fin trade, which made it difficult for most of them to open up to researchers. Face-to-face interviews were conducted with a total of 85 respondents, comprising 58 fishers and 27 traders in the three study communities. Interviews were conducted at the landing sites for fishers and mostly at the homes of traders. The interview lasted between 45 and 60 minutes per respondent. Interviews were conducted during the morning (08:00- 10:00), afternoon (12:00-13:00) and early evenings (16:00-18:00) in the landing sites and homes of respondents. The questionnaires were administered in local languages (Asante Twi, Fante, Nzima or Ahanta) by the first author, with assistance from local volunteers in their respective communities who served as interpreters when necessary, especially with the Nzima and Ahanta languages.

Data were collected with the permission from chief fishers (the person in charge of fish landing stations) and their elders (people who support chief fishers in deliberation and decisions taking at a particular fishing landing station) in their various landing communities. We preceded the interview by asking participants to give an oral consent to be interviewed. We informed every respondent of the purpose of the interview, the confidentiality of information provided, and the right to omit uncomfortable questions or withdraw from the interview at any stage, prior to the interview. Some respondents did not agree to be interviewed when we approached them. Five fishers immediately asked for permissions to withdraw from the interview when they were asked about shark trade even though they initially agreed to be interviewed. The information obtained from these respondents was expunged from the final analysis. We read questions from a semi-structured, standardized, questionnaire, which were identical across interviews and communities. Questions were mostly repeated and / or altered to ensure comprehension by respondents. Photo identification sheets were also used to confirm species names with respondents, when necessary.

The questions were used to elicit both qualitative and quantitative data on demographics, socio- economic attributes, livelihood strategies and fallback options from respondents. For the livelihood strategies, our pilot study revealed that both fishers and traders were unwilling to disclose the average income they earned from the livelihood stream they engaged in. We therefore asked respondents to state the average amount of income they obtained from their livelihood streams using a qualitative ranking scheme from 0–100% (categorized as; 0–20, 21– 40, 41–60, 61–80, 81–100). Subsistence consumption of shark products were recorded and categorized as often (i.e., once or more per week); sometimes (once per month); rarely (once or only a few times per year); and never (never at all). As a measure of fishers and traders well-being, we also captured information on their satisfaction with their work and income derived from shark fisheries. This was categorized as very satisfied, satisfied, dissatisfied, and very dissatisfied.

Questions were equally designed to collect data on main uses and sale prices of commercially important shark products (i.e., meat and fins), trade dynamics (including where the meat and fins are sold), and changes in the average prices of shark fins. The sales of shark fins are the sole responsibilities of the canoe owners (people who own canoe vessels) in their respective canoe business (Gelber, 2018; I. Seidu, pers. obs.). We therefore used the snowball sampling scheme to track down canoe owners and asked them about the shark fins and trade dynamics in their respective communities. The snowball sampling method, also known as referral or chain sampling, is used when potential participants are difficult to find (Newing, 2010). In this sampling scheme, research participants recruit other respondents for the study (Naderifar et al., 2017). Only 15 canoe owners (five from each community) participated in the interview, and provided information on the trade dynamics of shark fins. We specifically asked them to state the average price of eight commercially important and well-known shark species in their communities over a 15-year period. To facilitate the interview, four time periods were chosen together with events in which canoe owners were most likely to remember. These time periods were: i) 2005-2010, when fishers noticed a significant decline in sharks and pelagic fish species catch; ii) 2011-2013, where there was an embargo on the trade in shark fins; iii) 2014-2015, where the embargo on fin trade was lifted; and iv) 2018-2020, when data was collected. The average price for each shark species during each time period was used in the analysis.

In addition, we observed and collected catch and trade data on shark species in landing sites and used the data for our analysis on the changes in shark meat trade among the various communities. Shark trade data were collected during daylight hours at the three communities from March to June 2020. We recorded the species, size in cm (especially the precaudal length, as fins were often removed from the specimen), and the sale price of each specimen in Ghana Cedis (USD 1= GH¢ 5.77). When possible, we stood close by while fishers and traders negotiated on the prices of the specimen, or alternatively asked them about the prices after we recorded their sizes. A total of 397 shark specimens were sampled and their sizes recorded (Axim, *n* = 134; Dixcove *n* = 95; Shama *n* = 168). Sale prices were recorded for 713 specimens (Axim, *n* = 395; Dixcove *n* = 111; Shama *n* = 207).

### 2.4 Data analysis

All interview data were translated to English and were coded and analyzed using the Statistical Package for Social Sciences (SPSS) software, version 20. The Shapiro-Wilk test was used to test for normality in the data, prior to analysis (Zar, 2010). Chi-square contingency tests were used to test for significant associations between the relative income of the various livelihood streams among the study communities. The effect of gender, occupation, educational level, ethnicity and age group of respondents on relative income derived from shark fisheries and respondents‟ satisfaction with income from shark meat and their work were also tested using Chi-square contingency tests. The qualitative data were coded and analyzed using basic descriptive statistics in MS Excel spreadsheet and further presented in tables and figures.

The selling prices of sharks were mostly based on their sizes (primarily the precaudal length, after the fins had been removed), as no mechanism was put in place to weigh the specimen in the various landing sites. Thus, price per unit size (GH¢/cm) was computed for a total of 713 shark specimens in the three communities. Shark species that were less than 10 sale values were expunged from the analysis due to the low sample size. Prices at first sale and size data were tested to investigate if there were significant differences among the communities. Differences in price/cm of individual shark species among the three communities were tested with Kruskal Wallis tests and further compared with Bonferroni pairwise tests if there were significant differences among the communities. Statistical tests were conducted using PAST version 3.12 (Hammer et al., 2001), with a significance level of 5%.

## 3. Results

### 3.1 Socioeconomic characteristics of sampled respondents

The average household size was 3.59 ± 1.26 people (Table 2). Many of the respondents (*n* = 35) were between the ages of 36-45 years. Most respondents (81%) belong to the Fante ethnic group (Table 2). The ethnicity of respondents varies significantly among the study communities, with all respondents from Shama belonging to the Fante ethnic group. There was a significant difference in educational level of respondents among the communities, with most respondents having no formal education. The range of occupations of respondents also varies significantly among the study communities.

**Table 2.** Socioeconomic characteristics of respondents

| Socioeconomic characteristics | Axim<br>(*n* = 23) | Dixcove<br>(*n* = 22) | Shama<br>(*n* = 40) | Total<br>(*n* = 85) | *p*-value |
| --- | --- | --- | --- | --- | --- |
| Household size<br>(mean) | 3.78 (1.09) | 3.73 (1.12) | 3.40 (1.41) | 3.59 (1.26) | 0.478 |
| No. of dependent<br>household<br>members (mean) | 3.04 (1.30) | 2.73 (1.16) | 2.70 (1.32) | 2.80 (1.27) | 0.963 |
| Age of<br>respondents |  |  |  |  | 0.377 |
| 17 – 24 | 1 | 0 | 1 | 2 (2%) |  |
| 25 – 35 | 3 | 6 | 7 | 16 (19%) |  |
| 36 – 45 | 12 | 11 | 12 | 35 (41%) |  |
| 46 – 55 | 6 | 2 | 10 | 18 (21%) |  |
| 56 – 65 | 1 | 2 | 6 | 9 (11%) |  |
| 66 – 75 | 0 | 1 | 4 | 5 (6%) |  |
| Occupation |  |  |  |  | 0.020 |
| Fishery | 19 (22%) | 10 (12%) | 29 (34%) | 58 (68%) |  |
| Trading | 4 (5%) | 12 (14%) | 11 (13%) | 27 (32%) |  |
| Education level |  |  |  |  | 0.033 |
| Illiterate | 9 | 19 | 22 | 50 (59%) |  |
| Junior school | 10 | 1 | 13 | 24 (28%) |  |
| Senior High | 4 | 1 | 4 | 9 (11%) |  |
| Tertiary | 0 | 1 | 1 | 2 (2%) |  |
| Ethnic groups |  |  |  |  | 0.000 |
| Ahanta | 0 | 6 (7%) | 0 | 6 (7%) |  |
| Fante | 15 (18%) | 14 (16%) | 40 (47%) | 69 (81%) |  |
| Nzima | 8 (9%) | 2 (2%) | 0 | 10 (12%) |  |

1. Percentage sign (%) represents percentage of total respondents for each socio-economic variable; 2. Standard deviation in parenthesis without percentage sign (%); 3. Chi-square test for all statistical tests; 4. *p*-values represent significant differences in the socio-economic variables of respondents among the various study communities.

### 3.2. Fishing practices

All the study sites have both a multi-gear and multi-canoe fishery. The gears used by fishers are dependent on the size of canoe, fishing grounds (i.e., oceanic or coastal) and in many instances the finances of the canoe owners. The gear types predominantly used in these communities include longlines, handlines and trolling lines, purse seine nets, ring nets, drift gillnets, and bottomset gillnets. Sharks and rays are caught with two major gear types; drift gillnets complemented with longlines and bottom-set gillnets respectively. Baited hooks ranging from 110 to 250 are deployed as secondary longline gears, which are set alongside the drift gillnet in the same fishing grounds. These types of fishing gears are now widely used in the Axim, Dixcove and Shama communities and are used to target sharks and other pelagic species. Wooden canoes are the only vessels used by artisanal fishers in Ghana. Artisanal elasmobranch fishers use three types of canoe; large, medium and small (Table 3). Details of the fishing operations of the various types of canoe mostly used in the fishing communities and Ghana have been provided in table 3.

**Table 3.** Fishing operations and gears types used by the various type of canoes in the study communities and Ghana

| Fishing operations |  | Canoe types |  |
| --- | --- | --- | --- |
| Canoes used | Large | Medium | Small |
| Size of the canoe | 16-25 m long and 2-4 m wide | 9-15 m long and 1-2 m wide | 4-8 m long and 1-2 m wide |
| Outboard motor engine capacity | 40 HP | 15 HP, 25 HP, 30 HP or 40 HP | 8 HP, 25 HP or 40 HP |
| Gears mostly used | Drift gillnets | Drift gillnet, bottomset gillnets, and ring nets gears | bottomset gillnets |
| # of crews | 4-8 | 4-8 | 3-6 |
| # of fishing trips in a month | 4 times | 5-6 times | 6-7 times |
| Distance covered (km) | 129- 290 | 32-80 | 9- 16 |
| Trip duration | 6 days | 2-4 days | 1- 2 days |
| # of nets used | 20- 32 nets | 15- 20 nets | 15- 25 nets |

### 3.3 Reviews of Institutional frameworks on fishing and shark fishery in Ghana

Ghana has ratified a number of international and regional wildlife and fisheries frameworks that are relevant for the conservation of wildlife and fisheries resources. Regulations that apply to Ghana‟s marine megafauna are limited to dolphins and sea turtles. Regulatory action in Ghana is complicated by the socio-economic vulnerability of coastal fishing communities. However, the country has demonstrated its commitment to sustainable fishing as party to the 1982 United Nations Convention on the Law of the Sea (UNCLOS), the International Commission for the Conservation of Atlantic Tuna (ICCAT) and the Ministerial Conference on Fisheries Cooperation among African States Bordering the Atlantic Ocean (ATLAFCO). Ghana, as a member of the World Trade Organization (WTO), is subjected to the regulations governing fish trade and a signatory to a number of multilateral environment agreements including the 1973 Convention on International Trade in Endangered Species of Wild Fauna and Flora (CITES); the 1979 Convention on Migratory Species of Wild Animals (CMS) and the CMS Memorandum of Understanding on the Conservation of Migratory Sharks; and the 1992 Convention on Biological Diversity (CBD). Ghana is also a member of the Fisheries Committee for the West Central Gulf of Guinea (FCWCGG). The FWCGG is the regional body mandated to work towards a regional collaboration on management of the shared stocks and the regional integration of the national fisheries policies. Though the 1999 Food and Agricultural Organization International Plan of Action for the Conservation and Management of Sharks (IPOA-Sharks) is a voluntary and non-binding legal instrument, it was adopted to ensure the long term sustainable use of sharks by embracing the precautionary approach and calls upon maritime states to develop their tailored National Plan of Action for the Conservation and Management of Sharks (NPOA-Sharks). However, Ghana is yet to develop its own NPOA-Sharks.

Shark fisheries are not strictly regulated in Ghana. At the national level, the management of Ghanaian fisheries fall under the ambit of the Ministry of Fisheries and Aquaculture Development (MoFAD), which in turn delegates functions, including implementation, to a semi-autonomous body; the Fisheries Commission (FC). The FC was established under the Fisheries Commission Act of 2002 (Act, 625). The FC uses the Fisheries Regulation (L.I. 1968) and the Fisheries Law (PNDCL 256, 1991) to regulate the fishery sector in Ghana.

The Fisheries Act, 2002 (Act 625) is the main regulation of the fisheries sector of Ghana, which application is intended through the Fisheries Regulation, 2010 (L.1. 1968). The Act consolidates all the foregoing laws on fisheries, Decrees, Laws, Legislative instruments and other subsidiary/ subordinate legislation on the fisheries sector that are still in force. The Acts sets out to integrate international agreements into the country‟s national legislation. It sets out provisions for the regulation and management of fisheries, the development of the fishing industry, and the sustainable exploitation of fishery resources. Details of the laws and regulations relating to the management of fishery activities in Ghana are listed in Table 4.

**Table 4.**
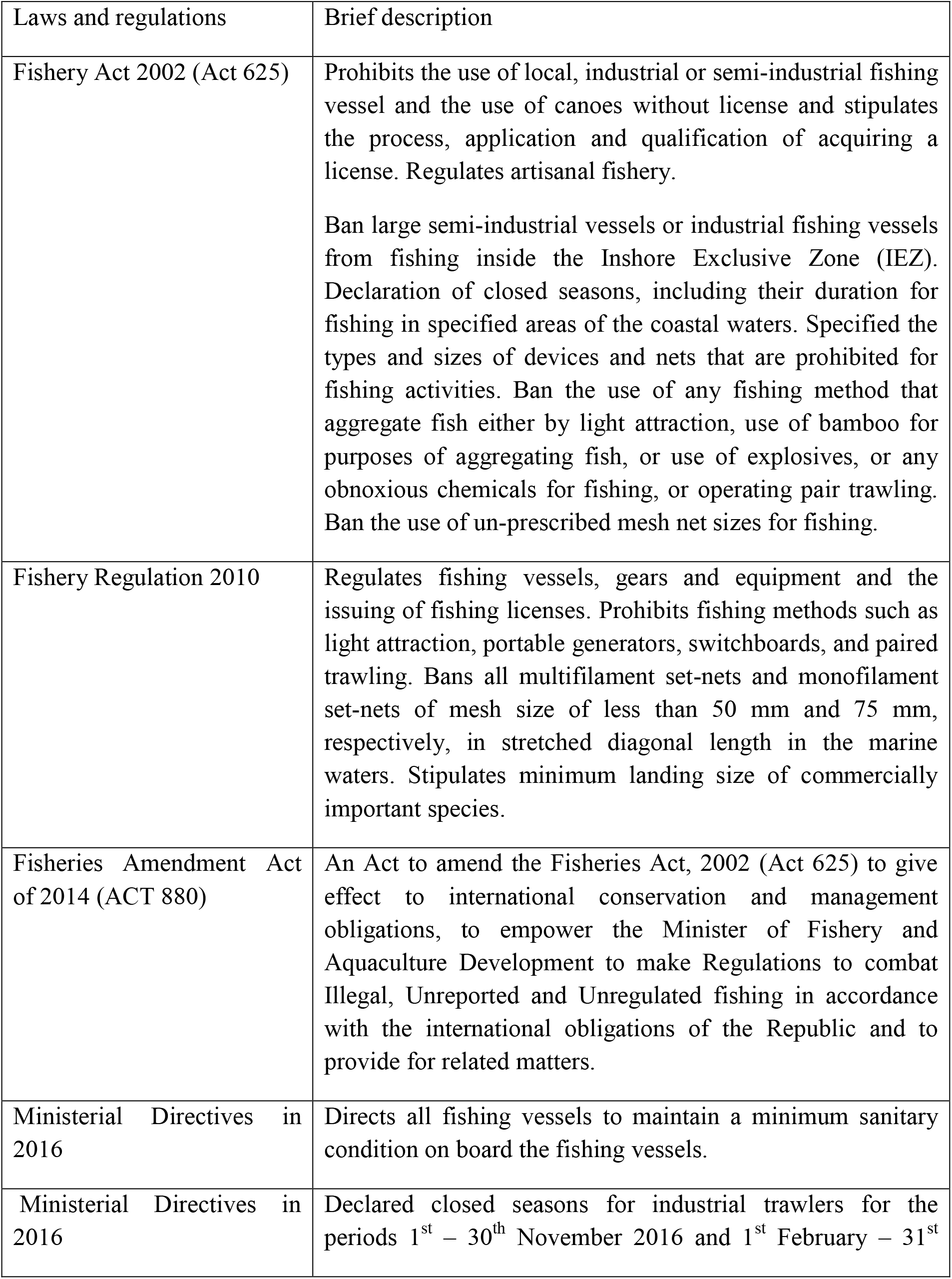
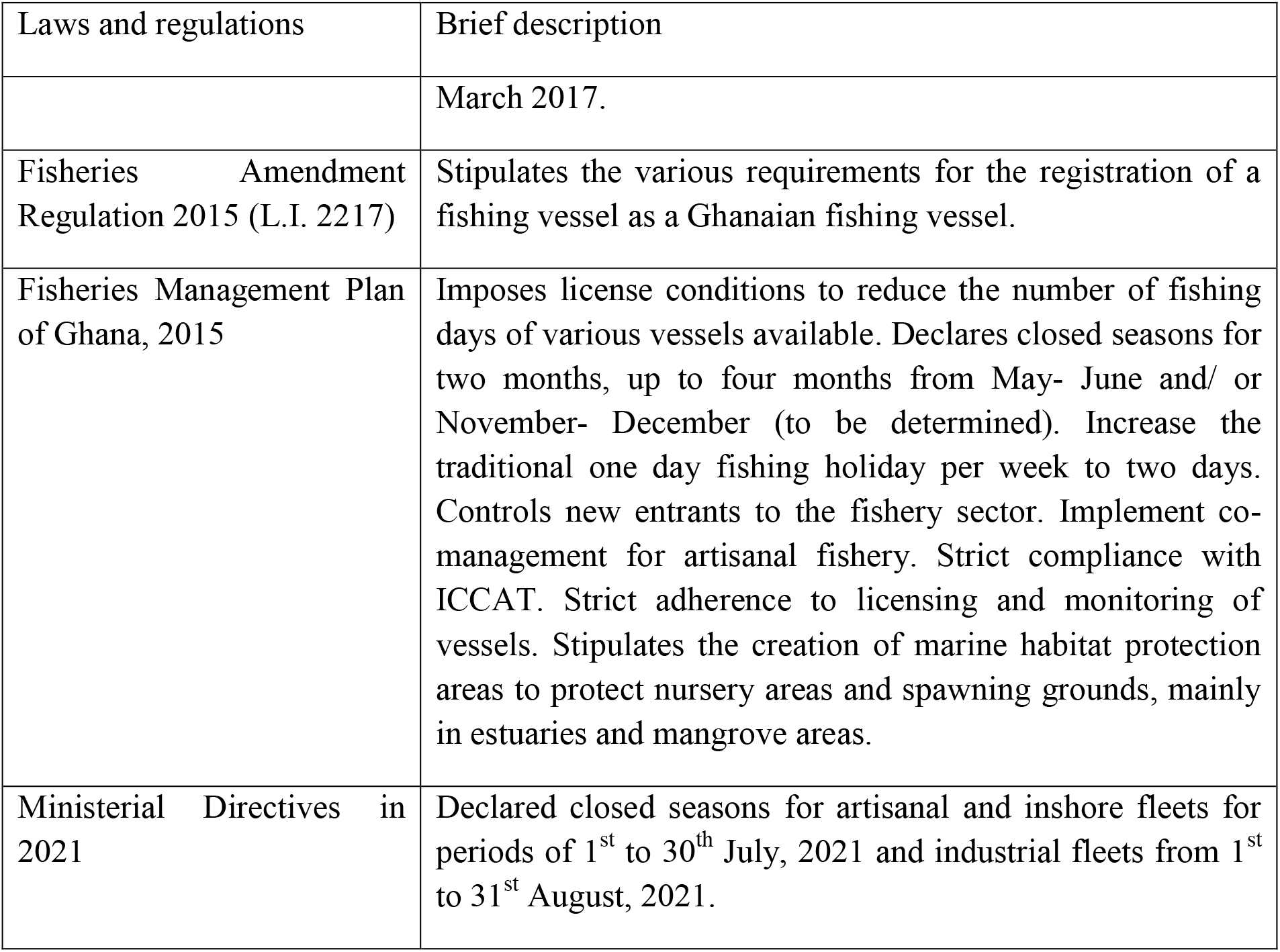
Laws and regulations governing Ghanaian fisheries

| Laws and regulations | Brief description |
| --- | --- |
| Fishery Act 2002 (Act 625) | Prohibits the use of local, industrial or semi-industrial fishing vessel and the use of canoes without license and stipulates the process, application and qualification of acquiring a license. Regulates artisanal fishery.<br>Ban large semi-industrial vessels or industrial fishing vessels from fishing inside the Inshore Exclusive Zone (IEZ). Declaration of closed seasons, including their duration for fishing in specified areas of the coastal waters. Specified the types and sizes of devices and nets that are prohibited for fishing activities. Ban the use of any fishing method that aggregate fish either by light attraction, use of bamboo for purposes of aggregating fish, or use of explosives, or any obnoxious chemicals for fishing, or operating pair trawling. Ban the use of un-prescribed mesh net sizes for fishing. |
| Fishery Regulation 2010 | Regulates fishing vessels, gears and equipment and the issuing of fishing licenses. Prohibits fishing methods such as light attraction, portable generators, switchboards, and paired trawling. Bans all multifilament set-nets and monofilament set-nets of mesh size of less than 50 mm and 75 mm, respectively, in stretched diagonal length in the marine waters. Stipulates minimum landing size of commercially important species. |
| Fisheries Amendment Act of 2014 (ACT 880) | An Act to amend the Fisheries Act, 2002 (Act 625) to give effect to international conservation and management obligations, to empower the Minister of Fishery and Aquaculture Development to make Regulations to combat Illegal, Unreported and Unregulated fishing in accordance with the international obligations of the Republic and to provide for related matters. |
| Ministerial Directives in 2016 | Directs all fishing vessels to maintain a minimum sanitary condition on board the fishing vessels. |
| Ministerial Directives in 2016 | Declared closed seasons for industrial trawlers for the periods 1<sup>st</sup> – 30<sup>th</sup> November 2016 and 1<sup>st</sup> February – 31<sup>st</sup> |
|  | March 2017. |
| Fisheries Amendment Regulation 2015 (L.I. 2217) | Stipulates the various requirements for the registration of a fishing vessel as a Ghanaian fishing vessel. |
| Fisheries Management Plan of Ghana, 2015 | Imposes license conditions to reduce the number of fishing days of various vessels available. Declares closed seasons for two months, up to four months from May- June and/ or November- December (to be determined). Increase the traditional one day fishing holiday per week to two days. Controls new entrants to the fishery sector. Implement co-management for artisanal fishery. Strict compliance with ICCAT. Strict adherence to licensing and monitoring of vessels. Stipulates the creation of marine habitat protection areas to protect nursery areas and spawning grounds, mainly in estuaries and mangrove areas. |
| Ministerial Directives in 2021 | Declared closed seasons for artisanal and inshore fleets for periods of 1<sup>st</sup> to 30<sup>th</sup> July, 2021 and industrial fleets from 1<sup>st</sup> to 31<sup>st</sup> August, 2021. |

Further, notwithstanding national law, fisheries activities are also under the responsibilities of traditional, customary authorities, like chief fishers and local fishing councils, who manage local fisheries and mediate conflict among fishers. The chief fishers and elders have their rules, taboos and norms, which support the regulation of fishing activities. Such taboos include prohibition of fishing on Tuesdays and the taboo on the catch and trade of whale sharks *Rhincodon typus* in some communities in Western Ghana.

### 3.4 Livelihood strategies

Fishing was the primary source of income and accounted for 58.5% of the total income of respondents (Table 5). Fish processing (16.0%) was the second most important livelihood strategy, followed by fish retailing (13.3%) and small businesses (2.5%). There were no significant differences in the mean income of all the livelihood strategies among the study communities, except fish processing (*X^2^ =* 34.44*, df =* 18*, p =* 0.011), which was higher in Dixcove compared to other communities (Table 5). In Axim and Dixcove, fish processing was the second most important livelihood activity after fishing. In contrast, fish retailing was the second most important source of income in Shama.

**Table 5.**
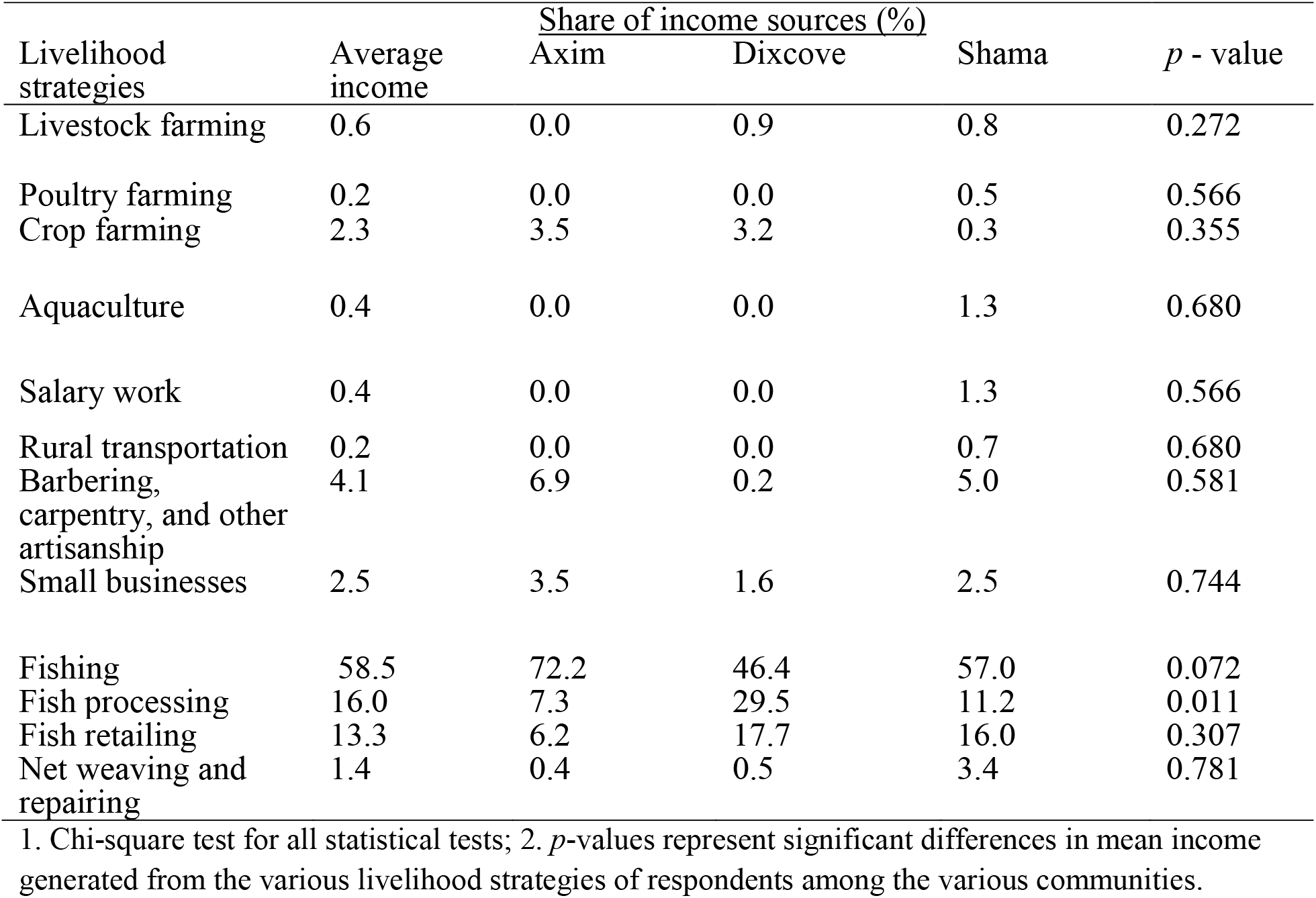
Mean income from livelihood strategies of respondents in the study communities

Across all communities, 67% of respondents only had one livelihood activity while 26% had two livelihood streams. Only 6% of respondents had more than two livelihood strategies even though they get less income from them compared to fishing-related activities.

### 3.5 Income from shark fisheries

Many fishers and traders generated between 80-100% of their income from shark fisheries. Most of the respondents in Axim (65% of 23 respondents) and Dixcove (68% of 22 respondents) received between 80-100% of their income from sharks. However, at Shama, only 35% of 40 respondents generated between 80-100% of their income from sharks. The remaining fishers and traders had income from sharks at different income ranges (Figure 3). Gender, occupation, educational level and ethnicity of respondents did not have any statistically significant influence on the range of income generated from shark fisheries. However, age of respondents had a significant effect on income produced from shark fisheries (*X^2^ =* 41.36*, df =* 20*, p =* 0.003), with 36-45-year-old respondents deriving more income from sharks than any other age group.

**Figure 3.**
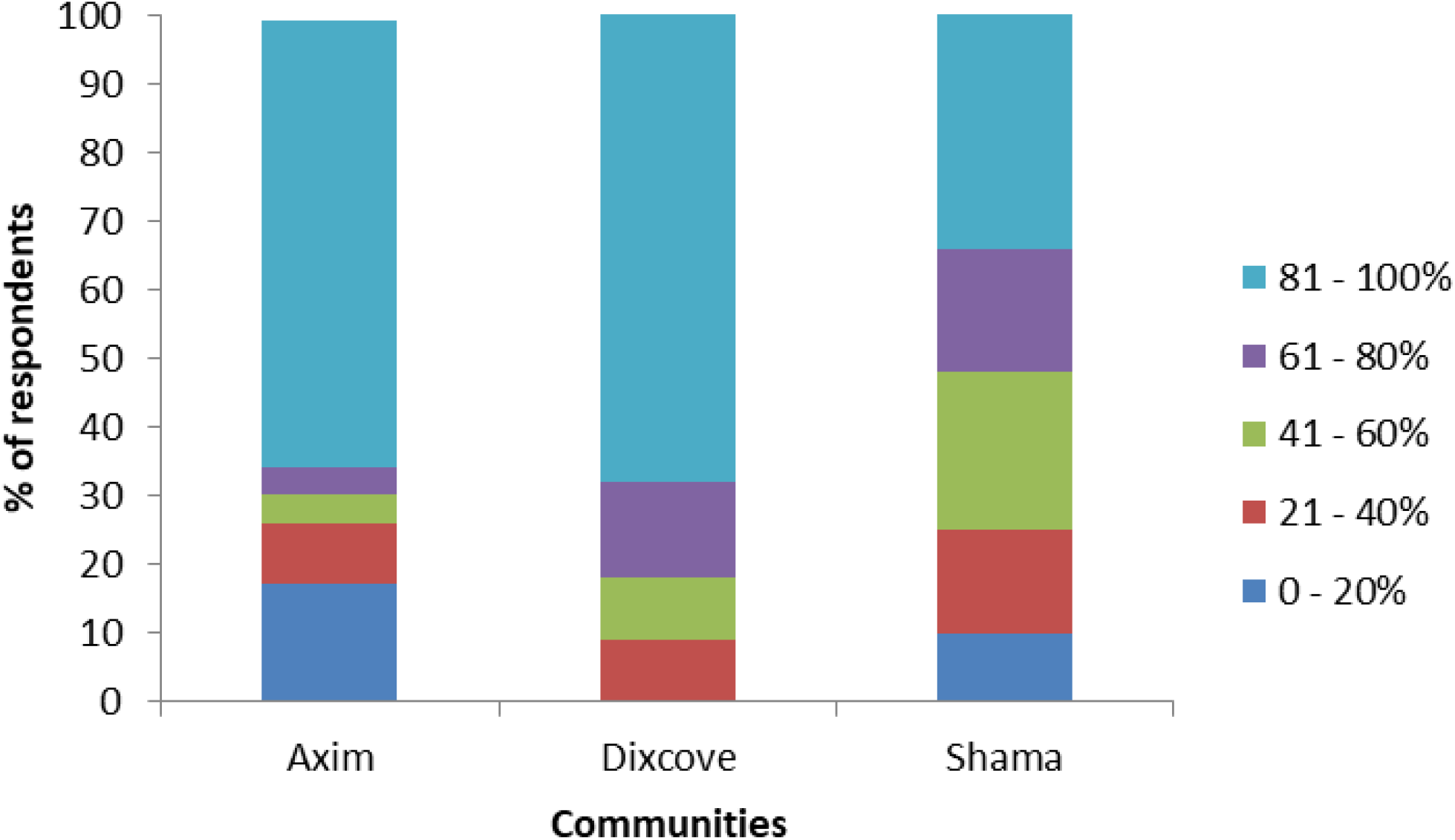
Income generated from shark fisheries in the study communities of Axim (*n* = 23), Dixcove (*n* = 22), and Shama (*n* = 40)

### 3.6 Subsistence consumption of shark meat

Consumption of shark meat was common in the three study communities (Figure 4). More than 80% of fishers and traders in each community ate sharks “often” or “sometimes”. Sixty-five percent of respondents at Shama stated that they often eat shark. Bull sharks (*Carcharhinus leucas*) and Hammerhead sharks (*Sphyrna* spp) were favored for consumption but other available species were also often eaten, including Blue Shark (*Prionace glauca*), Mako sharks (*Isurus* spp), and other requiem sharks (*Carcharhinus* spp). Many respondents (57%) in Axim ate shark “sometimes” and mostly preferred Sand Tiger Shark (*Carcharias taurus*) and Hammerhead sharks (*Sphyrna* spp). These fishers normally eat shark meat on Sundays and/or Tuesdays during fishing holidays and often ate Blue Shark and other requiem sharks, as they are easily available in their respective communities. At Dixcove and Shama, less than 10% of fishers and traders reported rarely eating sharks. Two of these fishers reported that they used to eat sharks often but now they no longer eat shark meat due to medical reasons, as their physicians have barred them from eating more meat. Only one trader had never eaten sharks before and attributed it to the nauseating way in which they are processed (i.e., the salting and/or smoking process).

**Figure 4.**
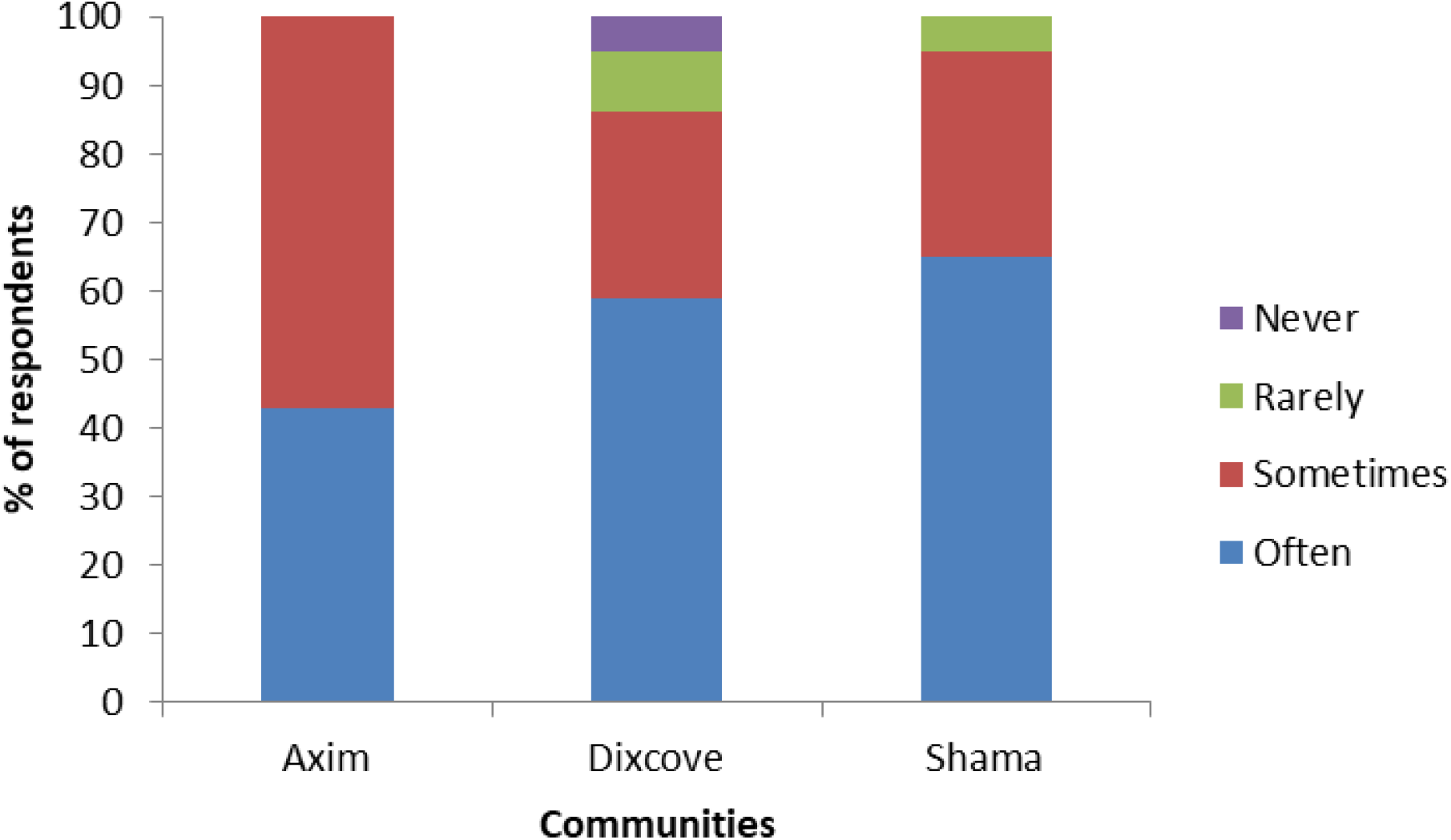
Frequency of subsistence consumption of sharks by fishers and traders. Stacked bars represent the consumption of sharks of the study communities in Axim (*n* = 23), Dixcove (*n* = 22), and Shama (*n* = 40)

### 3.7 Fallback livelihood options of fishers and traders

Fallback livelihood options are activities that fishers and traders would pursue to generate most of their income if there should be a ban on shark fishing and trading in Ghana. Non-fishing related livelihood streams were the major fallback activities both fishers and traders preferred to rely on if they could no longer engage in shark fisheries. Small businesses were the most common fallback livelihood option reported among the study communities (Figure 5). Some respondents, especially traders, mentioned some small businesses they would like to engage in as purchasing and selling of food items and other assortments in kiosks. Some fishers would like to acquire capital to purchase and sell clothing, men‟s underwear, shoes, and slippers in the capital city and other nearby markets. Transport business was the next important alternative livelihood option for some respondents, followed by artisanship like carpentry, barbering, and others. Some fishers were interested in driving taxis if they could get the skills in driving and money to purchase their own cars. Other respondents noted that they will be willing to work in the state transport cooperation (transport business own by the state) or other transport corporations in the country as cleaners, drivers, and conductors. Some respondents expressed their interest in exploiting and trading in other commercially important species like sardinellas, tunas, turtles, rays, anchovies, and cetaceans. They stated that they will modify their gears to target these species if there should be a moratorium on the harvest of sharks. Fewer than 20% of respondents from Dixcove and Shama stated that they will rely on government‟s support, aquaculture and poultry farming (Figure 5).

**Figure 5.**
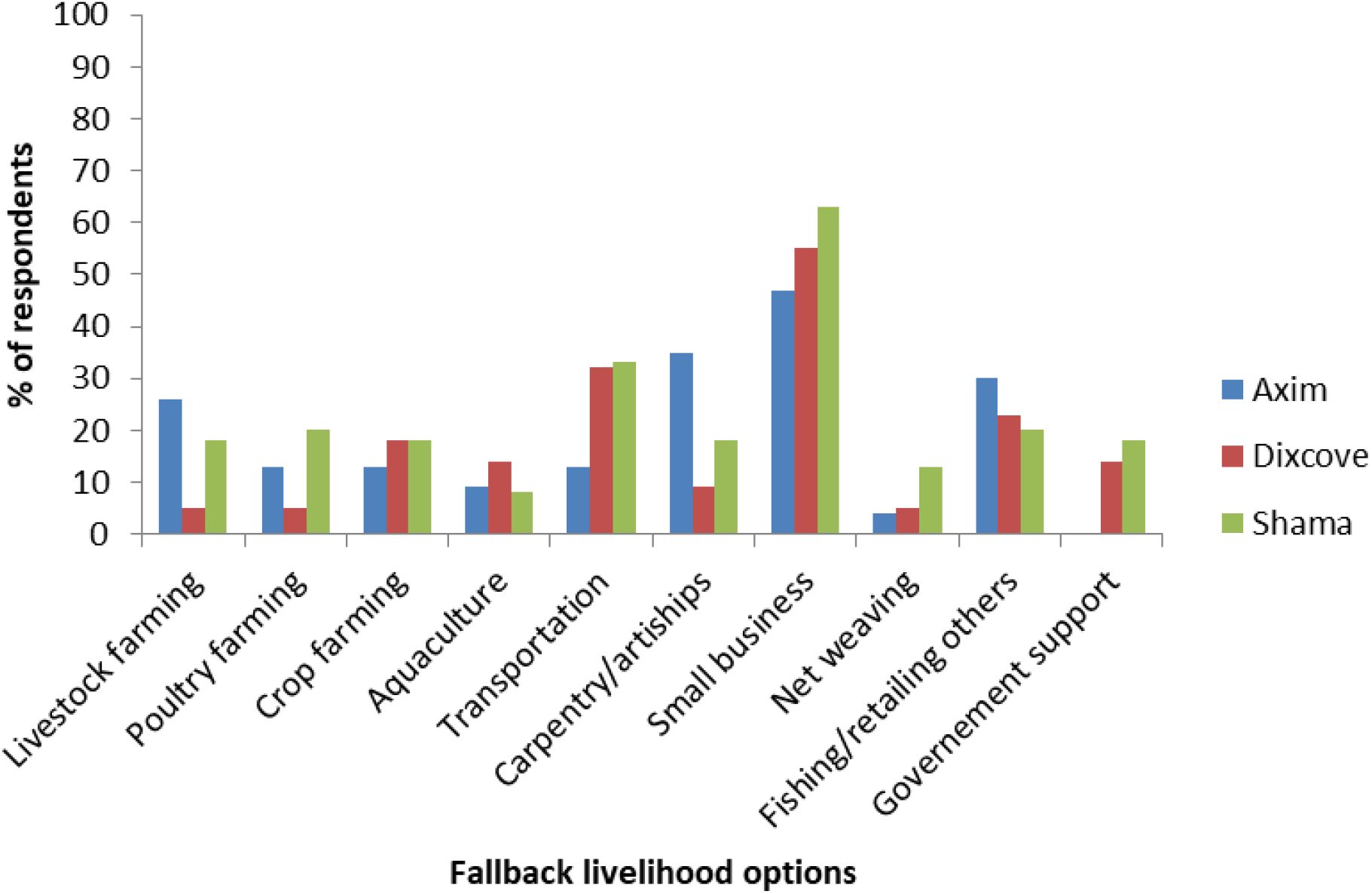
Preferred livelihood fallback options of fishers and traders if they could no longer harvest and sell sharks

When asked about the tools they require to implement these livelihood options, especially the non-fishing related activities, most respondents (68%, *n* = 58) stated funding as the major requirement they need before initiating these livelihood streams. Few respondents (35%, *n* = 30) also reported that they need to acquire the skills and technical know-how before they can kick-start their preferred livelihood options. However, five fishers who recounted having enough capital to start farming were increasingly concerned about the general unsuitable environment characterizing the sale of farm products and were afraid to lose their capital when they invest in farming.

### 3.8 Dynamics of trade in shark fins

A total of 15 respondents, who were mainly canoe owners, participated in the interview on shark fin trade as canoe owners are the ones solely responsible for the sale of fins in their respective canoe businesses. Fishers‟ perceptions on price dynamics over a 15-year period from 2005 to 2020 revealed that prices of all shark species had sharply reduced from 2011 to 2013 and increased 2-3-fold from 2014 to 2015 and continued to increase to 2018 (Table 6 and Figure 6). After 2018, however, prices remained virtually stable until 2020, as indicated by all canoe owners. The price of fins is mostly determined by buyers and marginally increases with time. Bull Shark, Hammerhead sharks, and Milk Shark (*Rhizoprionodon acutus*) consistently had the most valuable fins. Conversely, Blue Shark, Tiger Shark, and Thresher Shark are consistently the least valuable sharks in the local fin market. We initially anticipated some variation among respondents and communities but this was not the case as fishers indicated that prices of shark fins differ marginally among fishers and communities. Fishers reported that the local shark fin market is characterized by small number of foreign merchants who operate across all three study communities and therefore offer virtually similar prices.

**Figure 6.**
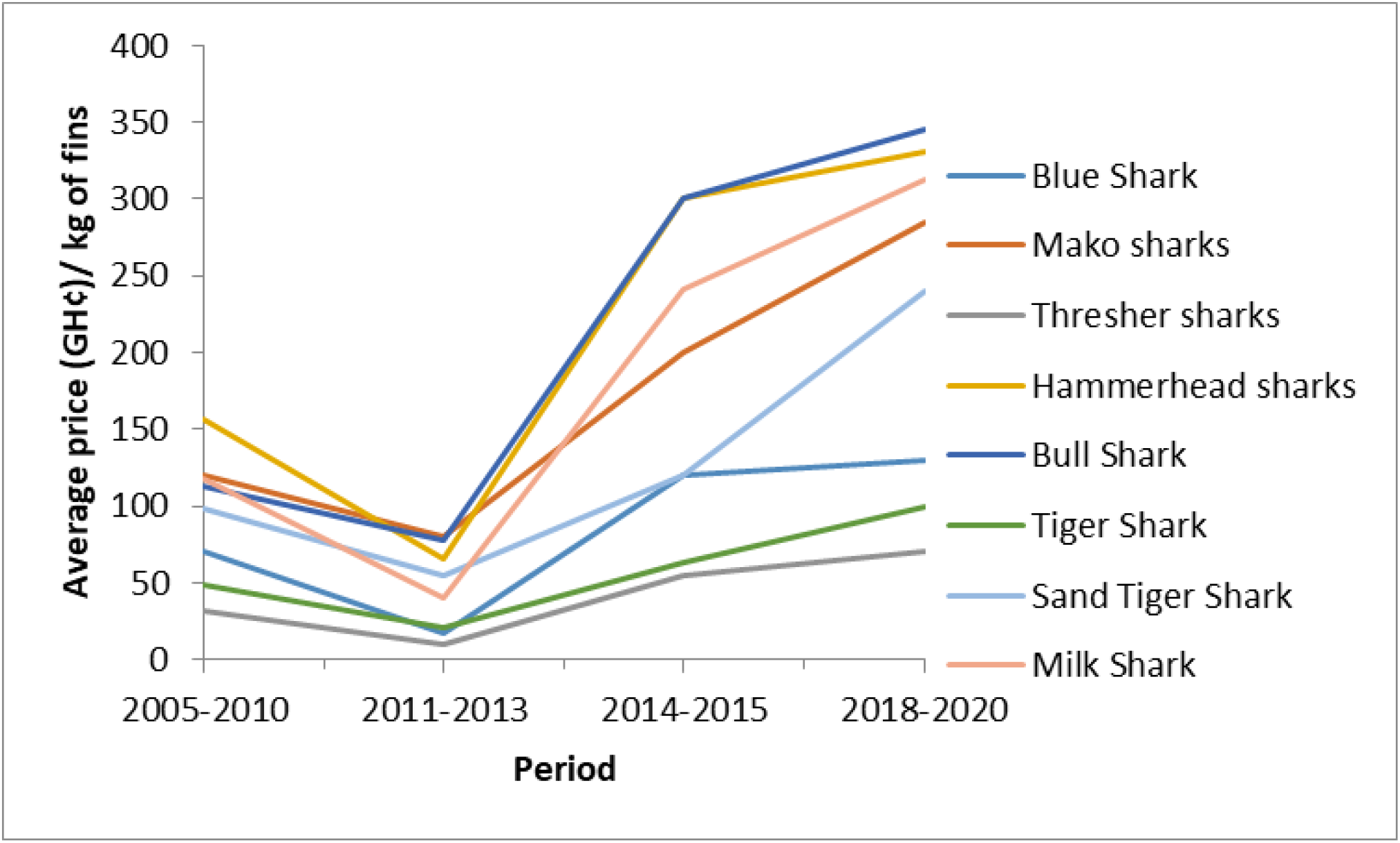
Trends in the prices for a kilogram of fins for eight shark species

**Table 6.**
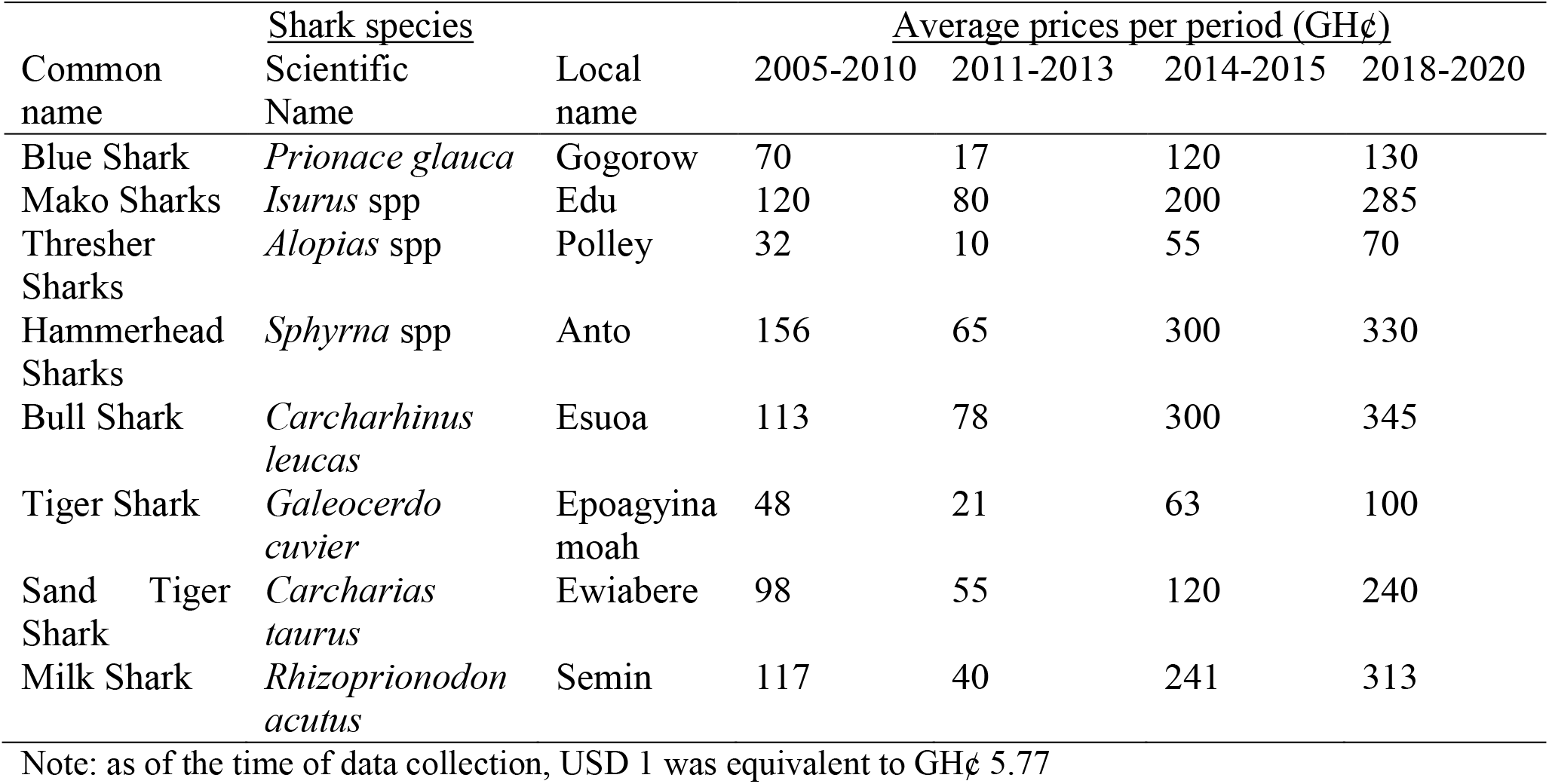
Average prices per dry kilogram of fins paid to canoe owners in the study communities

When asked about the details of buyers and factors that determine the price of shark fins, 67% of fishers recounted that they sold their fins to foreign merchants from Benin, Guinea, Mali and Senegal. Two canoe owners stated that they sell their fins to a Nigerian and has done so for the past six to eight years. Only one fisher from Axim claimed to sell his fins to a Chinese merchant since 2019. He indicated that the Chinese merchant used to conduct his business at Apam in the Central Region of Ghana but has recently relocated to Axim as the fin trade is booming there. Most foreign buyers do not stay within these shark fisheries communities, but rather they mostly stay on the outskirts of the various communities where canoe owners send their fins to them for sale. In certain circumstances, these foreign merchants go to the communities to buy the fins, especially when a canoe owner gets large quantities of fins and is unable to transport to the buyer‟s station.

Many fishers also stated that in rare situations, Ghanaian middlemen travel from one fishing community to another to buy fins from canoe owners. The prices they offer are relatively low compared to that of the foreign nationals. Fishers mentioned that they prefer to sell their fins to foreign nationals as they have built long standing relationships with them and they offer them good prices. Most fishers (87%, *n* = 13) reported that they mostly sell their fins from one to ten different buyers and have been doing so for over five to fifteen years now.

With the question of what the buyers do with the fins, 60% of fishers reported that they do not know who the buyers sell the fins to and/or what they do with it, and do not care to know because that is not their business. They stated that they were only interested in the money they receive from their sale and whatever the buyers do with their fins should not be their concern. Only three respondents stated that the buyers export the fins to China and other European countries and further said they are used for medications and as food in these countries.

Fishers reported that even though the prices are fixed some qualities are considered before the merchant buys the fins at that fixed price. For example, if the fins have not been properly sun- dried, the buyer will reduce the price on the fins. The dry weight and species type also affect the prices of fins. Fishers further recounted that seasons and level of demand for shark fins, especially in periods where there is scarcity of fins, affect the local market prices. Fishers were quick to add that scarcity of fins occur when there is shortage of premix fuel, which halts their fishing operations and in these periods if you get enough fins, you are likely to negotiate with the buyers for an increase in price. Further, ability of a buyer to finance fishing trips is also another factor reported to affect fins prices. Most foreign merchants sponsor the activities of shark fisheries by providing quick loans to canoe owners. The canoe owners pay back the loans after they get substantial shark catches, with the buyers paying a reduced price for the fins, as a form of deducting interest on the loans.

When queried about their satisfaction with the income they derived from the sale of fins, nine fishers stated that they are very satisfied or satisfied with the price and income they get from the sale of shark fins. These fishers recounted that the sale of shark fins gives them additional money to support their operations and stated that sharks are now increasingly becoming more lucrative than bony fish, as they get double income from the sale of fins and meat from sharks. Five fishers stated that they are dissatisfied with the price the buyers offer to them. Only one fisher reported that he is very disgruntled with the prices of fins as he feels the buyers are cheating them. He stated that he does not understand why for the past three years fin prices have not been increased, notwithstanding the increases in the price of fuel and the Ghana Cedis to the dollar exchange rate. He further stated that a merchant from Guinea once told him the prices of fins increases with the dollar exchange rate and sometimes with the fuel increment when he began selling fins, but later found that out not to be true.

### 3.10 Trade in shark meat

#### 3.10.1 Trade data from landing sites survey

Fins of medium- to large-sized sharks were removed, with the remainder auctioned in parts or whole, while smaller sharks were sold whole without the fins removed. After buying the specimen, traders would slice the meat into smaller pieces and then transport them to their various destinations. The prices at first sale of 713 specimens comprising nine shark species landed at the three study communities are documented in Table 7. Hammerhead sharks (*Sphyrna* spp) provided fishers with the highest mean price/length (cm), ranging from GH¢ 572/cm in Shama, to GH¢ 227.5/cm in Axim and GH¢ 216.7/cm in Dixcove. In Axim, Thresher sharks (*Alopias* spp) were the second highest valued species, averaging GH¢ 192.5/cm, followed by Mako sharks (*Isurus* spp) (GH¢ 175.7/cm) and Silky Shark (*Carcharhinus falciformis*) (GH¢ 176.4/cm). Conversely, Tiger Shark (*Galeocerdo cuvier*) (GH¢ 219.2/cm) was the second most valuable shark in Dixcove, followed by Bull Shark (*Carcharhinus leucas*) (GH¢ 147.5/cm), and Spinner Shark (*Carcharhinus brevipinna*) (GH¢ 105.0/cm). In Shama, Mako sharks (GH¢ 293.9/cm) exhibited the second highest mean price/cm, followed by Thresher sharks (GH¢ 200.0/cm), and Sand Tiger Shark (*Carcharias taurus*) (GH¢ 182.5/cm) (Table 7).

**Table 7.** Mean precaudal length (cm) and price (GH¢/cm) of nine shark species recorded in the three study communities in Western Ghana between 15 March and 11 June 2020

| Shark species |  | Mean length per community (cm) |  |  | Mean price per community (GH¢/cm) |  |  | *p* value |
| --- | --- | --- | --- | --- | --- | --- | --- | --- |
| Common name | Scientific name | Axim | Dixcove | Shama | Axim | Dixcove | Shama |  |
| Blue Shark | *Prionace glauca* | 174.4<br>(58) | 167.4<br>(46) | 176.6<br>(120) | 115.1<br>(150) | 148.2<br>(54) | 106.2<br>(144) | 2.066<br>E-10 |
| Silky Shark | *Carcharhinus falciformis* | 157.8<br>(6) | 129.6<br>(7) | 147.0<br>(6) | 176.4<br>(7) | 61.3<br>(8) | 96.0<br>(5) | 0.027 |
| Mako sharks | *Isurus* spp | 154.4<br>(27) | 132.0<br>(10) | 173.3<br>(14) | 175.7<br>(30) | 139.6<br>(11) | 293.9<br>(18) | 0.059 |
| Hammerhead sharks | *Sphyrna* spp | 161.0<br>(8) | 179.5<br>(4) | 213.8<br>(5) | 227.5<br>(6) | 216.7<br>(6) | 572.0<br>(5) | 0.126 |
| Tiger Shark | *Galeocerdo cuvier* | 155.0<br>(3) | 173.0<br>(4) | 211<br>(4) | 147.7<br>(7) | 219.2<br>(9) | 280.0<br>(7) | 0.172 |
| Thresher sharks | *Alopias superciliosus* | 170.0<br>(11) | 192.7<br>(6) | 228.6<br>(7) | 192.5<br>(12) | 138.8<br>(8) | 200.0<br>(10) | 0.538 |
| Bull Shark | *Carcharhinus leucas* | 143.9<br>(7) | 130.8<br>(6) | 106.7<br>(3) | 136.0<br>(5) | 147.5<br>(10) | 179.2<br>(6) | 0.562 |
| Spinner Shark | *Carcharhinus brevipinna* | 152.0<br>(9) | 115.3<br>(8) | 159.8<br>(5) | 51.8<br>(11) | 105.0<br>(10) | 108.0<br>(5) | 0.046 |
| Sand Tiger Shark | *Carcharias taurus* | 156.2<br>(5) | 158.3<br>(4) | 136.8<br>(4) | 56.0<br>(3) | 121.0<br>(5) | 182.5<br>(7) | 0.008 |

Note: 1. The number of specimens used to calculate mean values is reported in parentheses. 2. As of the time of data collection, USD 1 was equivalent to GH¢ 5.7; 3. *p*-values represent significant differences in mean price of shark specimen among the various communities.

The prices of shark meat were further analyzed to investigate if there were any statistically significant differences in the price at first sale among the various study communities. There was a statistically significant difference in the mean price/cm of Blue Shark (*Prionace glauca*) (*K*= 44.31, *p*= 2.070 E-14), Silky Shark (*K* = 7.12, *p*= 0.027), Spinner Shark (*K* = 6.13, *p* = 0.046) and Sand Tiger Shark (*K* = 9.64, *p* = 0.008) among the various communities (Table 6). Dixcove community differed significantly in the mean price/cm of Blue Shark in the Bonferroni pairwise comparison with Axim (*p* < 0.001) and Shama (*p* = 2.011 E-12). Similarly, the mean price/cm of Silky Shark differed significantly in the pairwise comparison between Axim and Dixcove (*p* = 0.020); the price of Spinner Shark varied significantly between Shama and Axim (*p* = 0.026); while Sand Tiger Shark varied between Axim and Dixcove (*p* = 0.012) as well as Shama (*p* = 0.019).

#### 3.10.2 Determinants of prices of shark meat

Generally, fishers reported that the price of shark meat is mostly dependent on the type of species and their sizes. Fishers stated that with equal sizes, Hammerhead sharks, Bull Shark and Mako sharks are the most valuable sharks, while Thresher sharks, Tiger Shark and Blue Shark are the least valuable in the local shark meat market. The level of demand and season were also other factors that affect the prices of shark meat. During traditional festive periods such as Kundum festival in Axim or Dixcove and Pra Nye-Eyi festival in Shama, fishers are mostly barred from going to the sea from one to two weeks and this affects the supply of shark meat in the local markets. Further, continuous shortages of premix fuel always halt the operations of fishers and thus, during these periods prices of shark meat are generally high for fishers who land sharks.

When asked about the changes in price of shark meat, most fishers and traders (62% of 85 respondents) indicated that they adjust prices of shark meat every time. Fishers confirmed that prices can either increase or decrease depending on their catch and the quantity a merchant buys from them. Only 29% of respondents stated that they change the prices of their meat every year or in season. Fishers also noted that the relationship with their buyers can affect the variations in price of their products. For instance, a trader noted that the prices of shark meat sold to local community members are less expensive than the prices they sell in other towns and regions in the country.

Most fishers sell their meat to two to five different merchants, with the highest bidder getting the product at the various landing sites. Only 31% of fishers sold their meat to a single merchant and in most cases such merchants happen to be their wives. In most cases, the wives provide a loan to support their husbands‟ fishing operations, which is repaid in kind by selling the meat directly to them. The merchant processed the meat in the form of smoking and salting and/or sun drying, which are sold in local markets as dried meat called “Kako”. Most traders stated that they sell their meat to local consumers in their various communities and outside their communities, especially in Takoradi, the regional capital, or Kumasi in the Ashanti Region, Tema and Accra in the Greater Accra Region, and Sefwi in the Western North Region of Ghana. Only four merchants recounted selling frozen shark meat to foreign nationals, mostly Chinese and Togolese, and have been doing that for more than two years. However, the traders were not aware whether these foreign nationals export the meat to other countries.

### 3.11 Satisfaction level of fishing and income from shark meat

As a measure of fishers and traders wellbeing in the shark fisheries, they were asked to indicate their satisfaction in the income they derived from shark meat and their work as a whole. Over 60% of fishers indicated that they are dissatisfied or very dissatisfied with the income they get from selling their meat (Figure 7a), while over 90% of traders were satisfied or very satisfied with their income derived from shark fisheries (Figure 7b). Dissatisfied fishers gave emotive responses to show how disgruntled they are with the prices traders offer to them. Many stated that catching sharks has become increasingly difficult and their work is now demanding more finances, energy and time and these are usually not taken into account when traders are buying their meat. In addition, 75% and 95% of fishers and traders respectively are satisfied or very satisfied with their work in fishing and trading shark meat. Most respondents reported that fishing-related livelihood activities are the only job they have learned and also their last livelihood resort to fall on for now, and that they are left with no option but to be okay with it. A trader stated that she learned how to process and trade shark and other bony fish since her childhood and that was the only training her parents gave her; but she is satisfied with her work because it at least offers her an income to pay her children‟s school fees and house rent.

**Figure 7.**
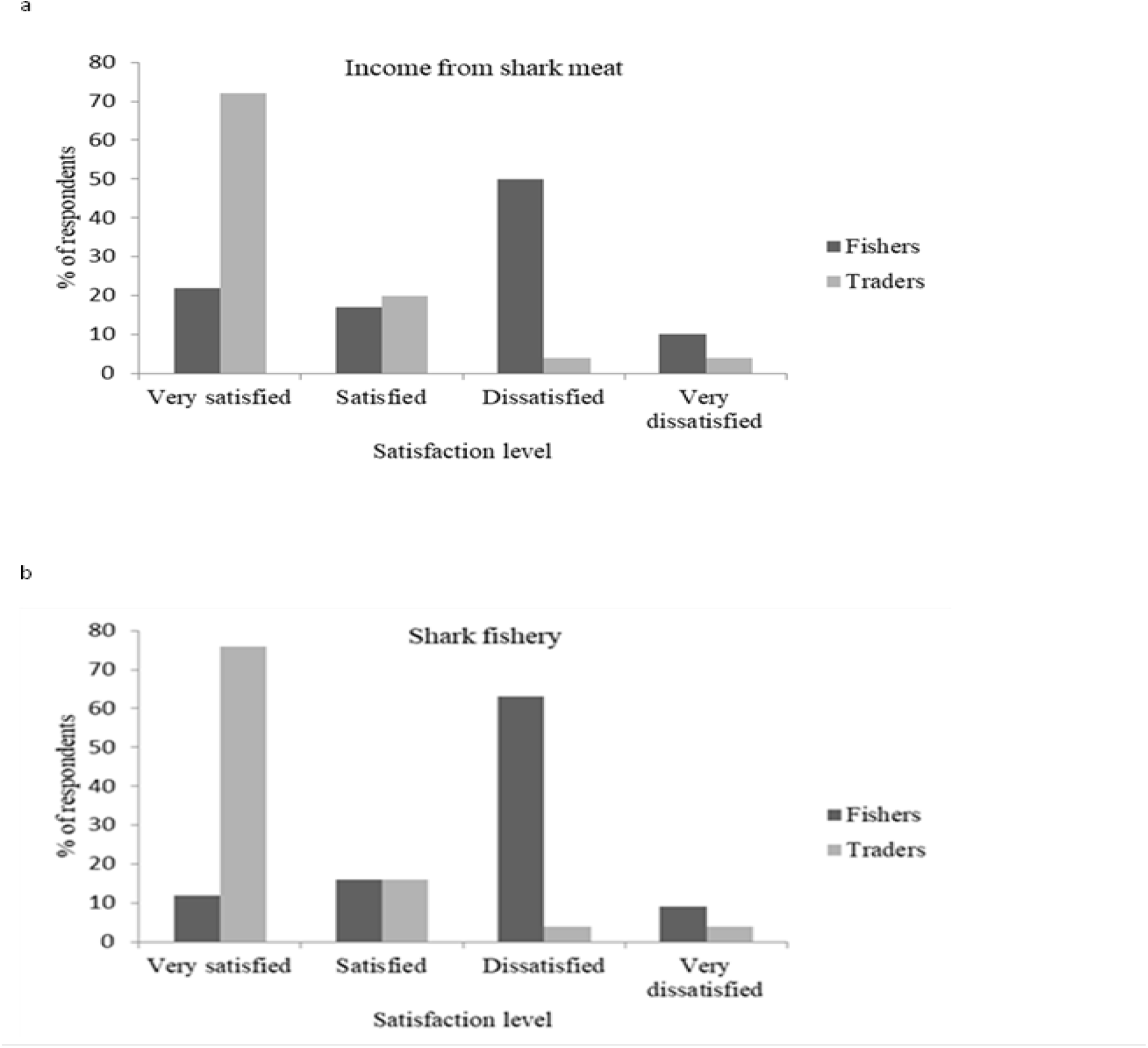
Fishers‟ and traders‟ satisfaction with income derived from fishing and selling shark meat

There was no difference in satisfaction with income derived from shark meat by educational level (*X^2^* = 14.84, df = 9, *p* = 0.095), ethnicity (*X ^2^*= 10.55, df = 6, *p* = 0.103) and age group (*X^2^* = 8.14, df = 15, *p* = 0.920). However, there was a significant effect of occupation on income satisfaction from shark meat (*X^2^* = 4.69, df = 3, *p* = 0.016), with traders exhibiting more satisfaction than fishers. Further, there were no significant differences in satisfaction with the work of fishing and trading fish between gender (*X^2^* = 1.88, df = 3, *p* = 0.059), or among occupation (*X^2^* = 2.19, df = 3, *p* = 0.053), educational level (*X^2^* = 9.21, df = 9, *p* = 0.418), ethnicity (*X^2^* = 5.67, df = 6, *p* = 0.461), and age group of respondents (*X ^2^*= 15.35, df = 15, *p* = 0.427).

## 4. Discussion

Our study is the first to characterize the livelihood strategies of shark fishers and traders and highlights the price dynamics of shark products in Ghana, and is one of the few from West Africa. Artisanal fishers in the study communities are considered to have limited livelihood opportunities. Most shark fishers and traders depend solely on fisheries-related livelihood strategies, while few had other alternative livelihood activities even though they generate less income from them. This finding corroborates the study by Barrowclift et al. (2017) regarding shark fishers and merchants in Zanzibar, where fishing was reported to be the primary occupation and main income source, with few fishers relying on secondary livelihood streams. In Ghana, Aseidu and Nunoo (2013) report that between 80% to 98% of fishers in Small London, Kpong, Ahwiam, and Elmina depend on fishing as their primary occupation and main source of income, while few (4-20%) had other minimal alternative livelihood options, including crop farming, livestock rearing, teaching, and trading in non-farm items. Sulu et al. (2015) analyzed a range of livelihood strategies adopted in the Malaita Province, Solomon Islands, and found that all respondents were engaged in multiple livelihoods activities, with fishing and gardening reported to be the most important livelihood streams. Generally, most available alternative income sources in the communities may entail unattractive returns on labor; a phenomenon that forces fishers and traders to expend almost all their time and energy on fishing-related strategies, which are deemed as lucrative in the study communities. Fishers and traders from Axim enjoy greater access to crop farming than those in Dixcove and Shama, owing to differences in soil and rainfall patterns (see Table 1); a reason why crop farming was among the secondary livelihood options in Axim community. Soil in Axim is fertile and combined with the high rainfall pattern favors farming activities in this area, which is among the mainstay of the people of Axim (GSS, 2014). Small-scale businesses and artisanship like barbering, masonry, hair dressing, carpentry, and others were secondary income sources for some respondents in Axim and Shama, likely related to having high levels of industrial development, which provide an enabling environment for such livelihood opportunities. Axim and Shama are capital towns in their respective districts, which are characterized with high population densities and significant levels of infrastructure development, with both private and government work opportunities (GSS, 2014). Prior to the survey, it was therefore expected that the relative infrastructure development in these communities will instigate many fishers and traders to rely partially on salary work for their alternative income source. However, only one fisher from Shama was a government worker and was receiving a monthly salary. This is as a result of high illiteracy rates of many fishers and traders, which disqualify them from applying for opportunities in government or private work that entails formal education. Even though many respondents did not have formal education, salary work was significant secondary income sources for fishers in other parts of Ghana (Asiedu & Nunoo, 2013), which contrast the findings of the present study. Further, in New Caledonia, salaried income work was an important secondary income source for fishers, and this was as a result of a high level of industrial development and a large mining sector in the country (Purcell et al., 2016).

Fishers and traders would mostly turn to non-fishing related livelihood activities, which include small business, transportation and artisanship as fallback livelihood options should there be a moratorium on shark fisheries. Restriction of fishers from shark fisheries may reduce fishing pressure, in the light of declining shark stocks (Ward-Paige et al., 2012). However, this study revealed that a significant number of fishers would simply switch to target other marine fish resources. The other marine resources stated by fishers such as turtles, rays, anchovies, and cetaceans are already threatened with extinction globally and require measures to safeguard them. This indicates that fisher‟s co-depend on various marine resources in small-scale fisheries because of the ease of shifting to other species, especially in light of marginal economic returns or restrictions on fishing certain stock (Purcell et al., 2016). Thus, a holistic approach needs to be adopted to simultaneously manage artisanal fisheries to encompass all economically important stocks, which may experience reduced fishing pressure, even when other stocks are the main target.

Similar to expectations that subsistence consumption of shark meat is prevalent in developing countries (Bornatowski et al., 2013), we found consumption common among fishers and traders in the study communities. Generally, the regular consumption of shark meat among fishers and traders suggest that shark meat represents substantial source of protein in the diets of the study communities and therefore over-exploitation of the shark stock may directly impact food security. In efforts to address food security, shark meat was promoted in the late 1950s as alternatives to augment the contemporaneous decline of bony fish, which has led to its wide utilization (Lehr, 2015). Currently, shark meat is widely traded and utilized as a cheap source of protein in many poorer communities in developing countries including Ghana (Bornatowski et al., 2013). Similar to the consumption pattern of shark meat in Ghana, Glaus et al., (2019) reported that Fiji‟s small-scale coastal shark fisheries are driven to mainly meet dietary needs. They reported that 79.3% of fishers that retain sharks utilized them as food source and/ or for cultural purposes and 19.8% sold shark products. Even in developed countries like Oman sharks are widely consumed and have formed the basis of many traditional food dishes (Henderson et al., 2006). Further, in the United Arab Emirates, fishers confirmed that the consumption of sharks has been integrated in their culture and has traditionally been consumed (Jabado et al., 2015).

Many fishers and traders generated between 80-100% of their income from shark fisheries, with most of these fishers from Axim and Dixcove. Shark fisheries are increasingly representing a key source of employment and providing major income for fishers in these communities. The high monetary incentive is the major driver of proliferation in shark exploitation in Ghana, as fishers are getting double their usual income in the form of sale of shark fins and meat. In agreement with these findings, Barrowclift et al. (2017) reported that most of fishers in Zanzibar that caught and sold elasmobranchs generated between 41-60% of their income from sharks, and 31% of merchants also got 61-80% of their income from selling elasmobranchs. In Ghana, Gelber (2018) found that the shark fin trade is the main income source for 80% of middlemen and 38% of canoe owners of the study population.

The fin prices of commercial shark species were found to be high between 2005 and 2010; the period where fishers reported that they experienced a dramatic decline in the catch of sharks and other large pelagic species. The decline in shark catch may have resulted in the high prices of fins, as demand might have been high in this period. Fishers started noticing a significant drop in fin prices in 2011, and the prices continuously remained lower until the end of 2013. Similar to our findings, fishers in Eastern Indonesia perceived changes in prices of shark fins over a 20 year period from 1992/93 to 2012/2013 and indicated that the prices steadily increased in 2002/2003, and decreased for all species in 2012/2013 (Jaiteh et al., 2017). Fishers in Eastern Indonesia gave diverse reasons for the fall in shark fin prices, including awareness campaigns targeting consumers in China, increasing demand for live reef fish at Chinese banquets, and international campaigns concerning the consumption of shark fin (Jaiteh et al., 2017), which contrast the reasons being given by shark fishers in Western Ghana. According to fishers in Ghana, from 2011 to 2013 there was an embargo on the trade in shark fins in Ghana, and merchants from neighboring West African countries migrated to their home countries. Several Ghanaian middlemen in these communities started buying and hoarding fins at very cheap prices during these periods. The moratorium on the trade on shark fins was thought to have occurred due to a number of reasons, including an investigation into the increasing cases of narcotics there were smuggled in fin cargo, of which the government of Ghana was concerned about (Gelber, 2018). Some canoe owners also related the ban on trade of shark fins to health issues, stating that there was spread of diseases owing to consumption of shark fins and this resulted in the ban on its exportation to China and other European countries. In 2014/2015, the fin trade ban was lifted and buyers started trading in shark fins and the prices increased exponentially as more buyers from neighboring West African countries moved into these communities to buy shark fins. In these periods, the demand for shark meat and fins was high but most fishing operations halted owing to long, incessant shortages of premix fuel. Only a few fishers were able to embark on long fishing trips to oceanic habitats and spent over six days at sea to catch sharks and these fishers got high prices for their fins. The prices of fins have since been increasing marginally from 2015 till 2017, and since 2018 the prices have virtually remained stable. In contrast to the current study, Glaus et al. (2019) documented a reduction in shark fin trade in 2017 in Fiji and linked the changes to the closure of the local sea cucumber market, which hampered the frequent visit of middlemen who used to encouraged shark targeting in the various fishing villages. Furthermore, annual shark fin income was estimated to have fallen by 75% following the sea cucumber fishery closure in that same year in Papua New Guinea (Vieira et al., 2017).

The prices of shark fins in Western Ghana were reported to vary among species. For example, the fins of Hammerhead sharks and Bull Shark were reported to be of high quality and therefore priced higher, while Tiger Shark and Thresher sharks were the least valuable species in the study communities. Similar to our findings, the lower caudal fins of Hammerhead sharks and Blue Shark and the fins of Shortfin Mako (*Isurus oxyrinchus*) have been cited to possess the best quality fin needles for human consumption by traders and regarded as among the most valuable fins in the international market (Clarke et al., 2007). Several factors influence the commercial value of shark fins globally, which include fin needles, type of fin (dorsal or caudal), the general appearance (thickness, color, length, and needle texture) and the species type (Clarke et al., 2007; Vannuccini, 1999) of which the latter is mostly known and used by fishers and merchants during fin trading in the study communities. Other factors reported by fishers as having an influence on the price of shark fins were dry weight, demand, and season. Though these factors were not statistically analyzed and inference on the prices was beyond the scope of this study, the data reported by fishers demonstrates the importance of fishers‟ knowledge in understanding the complex drivers influencing their fishing business and operations.

Further, the variation in prices of meat of Blue Shark, Silky Shark, Spinner Shark, and Sand Tiger Shark among the study communities may be attributed to the size differences of the specimen landed and sold. Sizes of shark species is an important factor that affects the prices of specimen, as larger specimens are given priority and priced higher. The mean size of Spinner Shark and Blue Shark was smaller in Dixcove while the mean size of Silky Shark and Tiger Shark were larger in Axim and Shama respectively, hence the variation in price among the communities (see Table 4). Additionally, the „quality‟ of the meat for consumption was reported to vary among species and therefore to influence the price of sharks as well. For example, species such as Mako sharks, Thresher sharks, and Hammerhead sharks are considered high quality and priced higher by fishers and traders, which concurs with the international shark meat market (Hanfee, 2001; Lehr, 2015; Rose, 1996). Similarly, fishers in the United Arab Emirates reported several species of sharks they considered most valuable, which included Hammerhead Sharks (Jabado et al., 2015). In contrast to the sale prices in Ghana, Bull shark were reported to fetch the highest price, resulting from their larger sizes in Zanzibar (Barrowclift et al., 2017). Further, Vannuccini (1999) reports that the spiny dogfish *Squalus acanthias,* sold in Italy for US$8.13– 9.91 per kg, was the most expensive shark species. Other interacting factors such as level of demand and season are also noted by fishers and traders to cause variation in shark prices in the study communities and this concurs with the study of Barrowclift et al. (2017).

Fishing marine resources contribute significantly to the degradation of the world‟s marine ecosystems and fishing pressure could possibly be reduced if primary actors, especially fishers in the industry, are induced to move out (Bavinck et al., 2012). Whether fishers may be inclined to do so or not, mainly depends partially on their wellbeing or satisfaction on their job and the profitability of their fallback options. Comparative studies have thrown light on the level of fishers‟ satisfaction on their professions. For example, Ruiz (2012) found that fishing is satisfying as an occupation, yet fishers can be dissatisfied about their earnings. Similar to our findings, most fishers in the present study were dissatisfied with the income earned from shark fisheries. This was because they expected a standard price for shark specimens but are mostly offered lower prices by traders. Fishers mostly compare their cost, time, and energy they expend on catching sharks and expect the prices to be higher than what they are offered by traders. The satisfaction of these primary actors in the shark fisheries industry is linked to the inadequate livelihood opportunities for fishers and traders in the study communities in, which they are invariably forced to stay and accept their occupation. Fishers have a high investment of their time in the fishery and often have few other viable livelihood options (Purcell et al., 2016). Most fishers may wish to switch job but the opportunities available are narrowed and most of them are not favorable to them owing to their training and level of education. The wellbeing of fishers, including their level of satisfaction, is advocated as an important consideration for development policy (Hair et al., 2016; Koczberski et al., 2006) and further offers more holistic means of assessing the social impacts of change in fisheries (Coulthard, 2012). Additionally, an insight of job satisfaction among the fishers and traders will support in developing management strategies that can offer required alternative work for these actors displaced by interventions for reductions in effort (Bavinck et al., 2012).

## 5. Conclusions and recommendations

Our study revealed that fishers and traders in shark fisheries in Western Ghana have marginal livelihood opportunities, with most respondents depending solely on fishing-related activities as their primary source of income. Secondly, shark fisheries contribute a significant income to these fishing communities, and shark meat is regularly used in the diet of both fishers and traders. Thirdly, non-marine fishing-related occupations, which include small business, transportation and artisanship, were the major fallback livelihood options both fishers and traders preferred to rely on if they are restricted from shark fisheries, but they require funding and adequate skills for their implementation. Fourthly, prices of shark fins reduced significantly between 2011 and 2013, but sharply increased in 2014/2015. The price of fins steadily increased until 2018, and has remained virtually stable till 2020. Hammerhead sharks *Sphyrna* spp have the most valuable fins and meat in the study communities. Fifth, over half of fishers were disgruntled with the income they get from selling shark meat, while most traders were satisfied with their income from sharks. Finally, most fishers and traders are limited with livelihood options and see fishing and trading of shark meat as their last safety-net and thus, are inclined to be satisfied with their jobs.

Inadequate alternative economic activities for fishers and traders of sharks may impede any management interventions to mitigate the impacts of their activities on shark populations. Thus, any management strategy would do well to consider various fallback livelihood streams outlined by fishers and traders. Failing to provide such incentives could result in opposition from fishers against any management intervention. However, the benefits of long-term higher sustainable yields would be worth the transition challenges.

## Funding

This work was supported by the Swiss Shark Foundation/ Hai-Stiftung, Save Our Seas Foundation and Flying Sharks.

## Acknowledgements

Special thanks go to the Zoological Society of London EDGE of Existence Program team members for their training, advice and mentoring. My heartfelt appreciation to Benard Seret for his immense contribution, mentoring, advice, and guidance for the species identifications and towards the successful completion of the study. Finally, the authors are grateful to Bukari Saphianu, Paul Tehoda, Emmanuel Amoah, Clement Sullibie Saagulo Naabeh and Adomako Ohene as well as the local volunteers Isaac Assefuah, Kingsford Osei, Jona, Timothy Amiah and Ben Adjei for their role in field data gathering. We are grateful to the staff of the Western Regional Fisheries Commission, chiefs, Chief fishers and their elders, fishers, traders and people of the five study communities for their cooperation and support in making this study possible.

## Notes

### Competing Interest Statement

The authors have declared no competing interest.

